# Introgressive Descent and Hypersexuality Drive The Evolution Of Sexual Parasitism and Morphological Reduction In a Fungal Species Complex

**DOI:** 10.1101/2023.01.10.523206

**Authors:** Fernando Fernández-Mendoza, Eva Strasser, Ivan Frolov, Jan Vondrák, Lucia Muggia, Helmut Mayrhofer, Ester Gaya, Martin Grube

## Abstract

Taxonomists consider species as discrete units of biological organization, which are subject to a continuous process of evolutionary change and are connected through their shared ancestry. However, the continuous nature of evolutionary change is difficult to reconcile with the discrete outcome of speciation, especially where species boundaries are permeable. A good example of this inconsistency is the lichen genus *Pyrenodesmia*, which shows a high morphologic and genetic diversity that that defies systematization by taxonomic or phylogenetic methods. Here we show that hybridization explains the presence of discordant morphs and that European species are interconnected through cross-mating in a single reproductive network, a syngameon, despite which species remain largely distinct and distinguishable. Whole genome data reflect the important role played by genome defense mechanisms in the genomic stabilization of fungal hybrids. The recurrence of Repeat Induced Point mutations (RIP) shapes genomes with islands of suppressed recombination and loss of gene content, which in turn generates a feedback loop reinforcing the lack of reproductive isolation through the loss of heterokaryon incompatibility and a tendency towards morphological reduction.

## BODY

The idea that life is structured into discrete evolutionary units (i.e. species) is central to biological science, and articulates the way we understand, organize and communicate biodiversity. Considering species as the natural unit of biological organization, evolution, and ecological interaction is a widely accepted simplification that allows linking evolutionary biology with all other disciplines in biology, from ecology to molecular biology, but overlooks significant aspects of the way organisms evolve. First and foremost, species cannot be considered as the only unit of selection and evolution because both processes are acting simultaneously across all levels of organization (*1*–*3*). Second, there is no unifying consensus on what species are, their necessary properties (*4*–*15*), and how they integrate into a continuous process of evolutionary change. This has led to discussing whether species provide any usable insight when dealing with prokaryotic lineages (*16*–*19*).

The need to consider evolution and selection as multilevel processes has become ever more apparent with the increased availability of sequenced genomes. Horizontal transfer of genetic information between distant organisms is more frequent than once thought and has been identified as a key element in habitat adaptation (*20*), which reinforces the idea that communities function as supra-specific evolutionary units (*21*) shaped by the transfer of information through non-genealogical bonds (*3*). Similarly, the study of hybridization between closely related plant and animal species has long identified species complexes in which phenotypically differentiated species are interconnected through gene flow. Both syngamea (*22*, *23*) and metaspecies (*24*, *25*) can be interpreted as supra-specific units of evolution. Conversely, the detailed comparative study of genomes identified significant asymmetries in the distribution of recombination, hybridization and mutation among genomic regions (*26*–*28*). This heterogeneity, which influences and is influenced by the underlying genomic architecture, results in identifying genomic islands of speciation (*29*) and local adaptation (*30*), which can be interpreted as infra-individual units of selection.

Next, the use of molecular genetic characters and phylogenetic methods (*31*) in fungal research fundamentally changed the way diversity is studied and understood. In the first instance, genetic characters allow identifying cohesive and diagnosable (*6*) groups of specimens even when the “discontinuity of organic variation”(*32*) is not observable using morphology and chemistry. Furthermore, the presumed objectivity of genetic characters and phylogenetic methods resulted in a widespread redefinition of fungal taxa in terms of monophyly (*33*–*37*) and a widespread identification of cryptic species (*38*–*51*) in fungi. This resulted in considering that the “true diversity of fungi” (*52*) had been obscured by overly wide taxonomic concepts based on morphology. To open the treasure chest of fungal diversity, molecular taxonomists progressively incorporated growingly complex methods of unsupervised species discovery and validation (*53*–*61*). However, recent surveys tone down notably the former claims made on the extent of fungal diversity (*62*) former claims on hidden fungal diversity. In addition to the known limitations of DNA barcoding when used for species delimitation, validation, and recognition (*63*), phylogenetic methods of species delimitation (*31*), despite their progressive refinement (*64*–*71*), are quite limited in their application and share the same caveats. First, they cannot accommodate gene flow (*72*–*74*) unless species limits are imposed a priori (*75*–*83*). Second, they are only meaningful when populations don’t deviate significantly from classical population genetics models (*84*). Both limitations have been systematically overlooked despite the growing evidence that species boundaries are semi-permeable (*85*) in lichenized (*86*–*88*) and non-lichenized fungi (*89*–*91*), and the evidence that the outcome of reproduction may not adhere to mendelian inheritance (*92*–*94*) or the simplistic expectations of Hardy-Weinberg equilibrium.

Here we evaluate the diversity observed in the lichen genus *Pyrenodesmia* from multiple points of view to quantify the usability of phylogenetic species concepts and the extent to which hybridization and reticulation shape its evolutionary history. Using a multilevel approach we evaluate the presence of supra-specific evolutionary units, how they influence the reproductive and evolutionary strategies of species within the genus, as well as the influence that hybridization may have on genome architecture and mosaicism.

We ask the following questions: (i) What is the diversity of the genus *Pyrenodesmia* in the European continent? (ii) Are phylogenetic species concepts usable? How many phylogenetic species are there? (iii) Is there evidence of hybridization in the genus? (iv) What are the reproductive units (v) What are the genomic consequences of reticulate evolution and hybridization?

### One-third of European *Pyrenodesmia* specimens cannot be identified using existing morphological species concepts

The genus *Pyrenodesmia* s.s. (*34*) is a widespread lineage of lichenized fungi within the family *Teloschistaceae* whose species form mostly saxicolous crusts and share the lack of anthraquinone pigments as the most obvious, albeit paraphyletic character (*95*). Its species are quite diverse in apothecial morphology, thallus pigmentation, and thallus development, ranging from the large crusts of *P. transcaspica* to the endolithic thalli of *P. erodens, P. albopruinosa*, and *P. alociza* (Fig 1a). Despite the substantial morphological and ecological differences, the taxonomy of the genus remains unclear. The main culprits are the low signal found in micromorphological characters and the abundance of intermediate forms which are difficult to interpret. As a result, pre-molecular taxonomic works (*96*–*98*) used a broad concept of species (i.e. *P. variabilis* or *P. circumalbata*) and their classifications were often disparate. Meanwhile, recent phylogenetic surveys consistently describe new species using monophyly as diagnostic criterion (*99*–*106*), suggesting that a significant proportion of the species diversity remains unidentified.

**Fig. 1.**
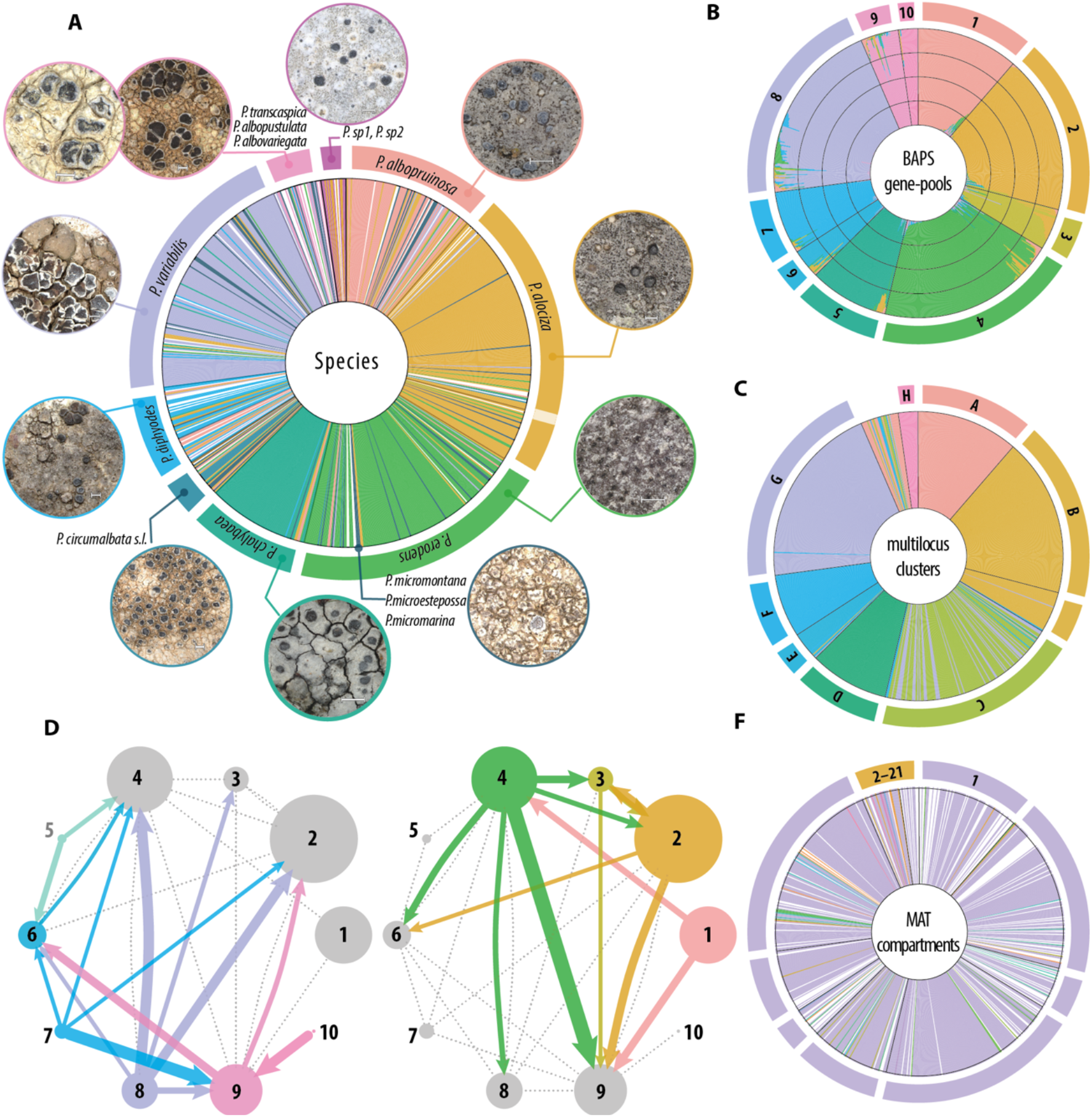
Taxonomic, phylogenetic, and population summary of the dataset. Figures a-c and f provide a graphical summary of the structure of the dataset using the assignment of specimens to gene-pools estimated in BAPS as organizing scaffold. **a**| Assignment of individual specimens to the twelve most relevant morphospecies. A more complete taxonomic profile is provided in the supplementary material. Each color signifies the assignment to a species or species group. The scale bars in all habitus pictures amount to 500μm. **b**| Assignment to admixed gene-pools (populations) obtained in BAPS, each specimen is represented by a stacked barplot. The color highlights the proportion of the variability assigned to each of the ten estimated admixed pools. **c**| Assignment to putative species (coassignment clusters) using a multilocus interpolation of single locus bGMYC assignments. All clustering algorythms consistently identified eight clusters. **d-e**|Estimates of migration between the BAPS genepools calculated using in migrate-n. For clarity, the network is split into two subnetworks highlighting gene flow from pools mostly containing well-developed epilithic species and those from pools comprising mostly reduced endolithic species. Each node is a numbered gene pool, correlative to the numbering used in fig.1a. the size of each bubble represents the estimated mutationally scaled population size (θ), and the edges represent immigration estimates recalculated to represent the number of immigrants per generation (2Ne). The size of each edge represents the migrant estimates provided in the supplement. The network was simplified to include only strong connections (2Ne>2). **f**| Assignment to Mating compartments using the unipartite network of MAT idiomorph cooccurrence. The network contains a main subnetwork comprising most specimens and 20 secondary ones.

As starting point we opted for re-evaluating the species diversity of the genus *Pyrenodesmia* in a well-studied region, so we assembled a dataset comprising mostly saxicolous species growing on limestone across Southern Europe (Table S1, Figure S1). Based on the taxonomic literature we expected to find at least seventeen distinct species, six of which are common in Southwestern Europe (*P. albopruinosa, P. alociza, P. chalybaea, P. circumalbata, P. erodens* and *P. variabilis*), five are difficult to differentiate from each other (*P. diphyodes, P. helygeoides*, and *P. microstepossa, P. micromarina* and *P. micromontana*) and one is rare (*P. badioreagens*). The remaining five species occur outside of the studied region (*P. albopustulata, P. albovariegata, P. concreticola, P. molariformis* and *P. transcaspica*) and were included to provide a wider geographic and taxonomic context. The morphological study resulted in identifying 42 operational units (species and pseudospecies) of which twelve were collected outside of the main study area, nine are clearly identifiable, three have been described based on molecular characters and are doubtful (Figure 1a) and the remaining eighteen show intermediate character combinations and are difficult to place, being potentially undescribed species.

The consensus in lichenology is that molecular taxonomic methods are necessary to overcome the bias introduced by overly simple and subjective morphological species concepts (*52*, *62*). To address diversity, most surveys follow a similar recipe (*63*): assembling a phylogenetic dataset, proposing a set of operational taxonomic units (OTUs), which are finally validated using a multilocus framework, often based on multispecies coalescent (*107*–*110*) or an analogous strategy (*111*). To evaluate the species diversity of *Pyrenodesmia*, we assembled a phylogenetic dataset including 824 lichen specimens and five nuclear loci (Tables S2-S4). The need to phase sequences from specimens that included dikaryotic mycelia resulted in a 0.98 complete data matrix of 910 phased genotypes (Table S3).

The resulting dataset showed very high haplotype and nucleotide diversity (Table S3). Haplotype diversity is reflected in low levels of sequence duplication, with 0.40–0.56 of sequences being unique, meanwhile the high nucleotide divergence between sequences is best reflected by having between 0.38 and 0.46 of sites being parsimony informative across all nucleotide alignments, excluding the outgroups.

This is very different to what we encountered in previous macrolichen surveys (*112*– *117*) where species tend to comprise: a) few closely related haplotypes, identifiable as discrete lineages in phylogenetic trees and network reconstructions and b) few overrepresented haplotypes common across large geographic regions (*115*, *118*, *119*). The observed phylogenetic continuum is similar to that of *Lecanora polytropa*, a species complex proposed to comprise 70 putative species worldwide (*62*).

### Interpreting diverging genotypes as separate species artificially inflates the estimates of species diversity

An immediate consequence of interpreting highly diverse single locus alignments in terms of species delimitation is the proposal of high numbers of putative species (Table S3). Widespread, albeit simplistic distance-based (ABGD and ASAP) and phylogenetic methods (GMYC) identify an average of 180 species, although the number of proposed entities has a wide range between 23 and 368 OTUs depending on the locus, method, and threshold used for delimitation (Table S3). The high nucleotide divergence found between haplotypes results in a high abundance of singleton clusters (0.90–0.63), comprising a single observation, reflected in the strong numerical discordance between the number of identified clusters and entities in GMYC. The multitree implementation of the GMYC algorithm (bGMYC), aimed at integrating phylogenetic uncertainty often results in the most conservative estimates, although they also depend on the criterion used to obtain a consensus from the coassignment matrix: the average silhouette width of k-medoids estimated 15–97 OTUs, the median number of partitions 5–50 OTUs.

In general terms, the main OTUs statistically associate with the identified morphospecies across all loci, the correspondence between them is never complete, and specimens with very different morphology tend to be clumped together (Table S3, Supplementary material). Furthermore, the OTU delimitations are not fully concordant between loci. To integrate all sequenced loci on a single delimitation strategy we combined the bGMYC coassignment matrices obtained for single loci into a single dissimilarity matrix, which was used to estimate a consensus delimitation using k-medoids clustering and average silhouette width as criterium (Figure 1c). This consensus delimitation consistently partitioned the dataset into 8 genetic clusters, which are largely coherent with single-locus OTUs (Cramer’s V = 0.81–0.97, Supplementary information), morphology-based identifications (Figure 1a, Cramer’s V = 0.77 Supplementary information), and estimates of population structure (Figure 1b, Cramer’s V = 0.932). This multilocus consensus, limited as it is by its mathematical simplicity, highlights the inadequacy of using single locus datasets to generate species hypotheses and of interpreting speciation as the only source of genetic variability.

### Phylogenetic analyses suggest the presence of widespread hybridization

The limitations of phylogenetic methods of species delimitation are well known (*31*) and have been thoroughly discussed in the fungal literature (*52*), despite which they remain useful for species discovery (*62*, *63*). Most models used in species delimitation rely on strong assumptions about the contribution that reproductive isolation, genetic drift, and mutation have to the genetic variability at macro and microevolutionary levels. Most importantly, phylogenetic species concepts rely on considering species as reproductively isolated lineages, a condition that is not always met in natural populations.

In *Pyrenodesmia*, the phylogenetic signal in the identifiable phenotype-based species and the concordance between single-locus phylogenies are high and statistically supported, but still partial. The average concordance between loci, estimated using Nye’s similarity metric (*120*) on Bayesian tree distributions, amounts to 0.45 (0.15– 0.70) of the average concordance found within loci. The concordant fraction within loci averages 0.34–0.42, this is significantly higher (ML t = 34.99, p-value<2.2e-16; By t = 245.11, p-value<2.2e-16) than the random expectation (ML 0.15, By 0.19), and about half of the concordance observed within loci (By 0.69 t = −100.79, p-value < 2.2e-16), which provides more reasonable null expectation. Despite the concordance found between loci, the consensus phylogenetic network reflects a strong incongruence between gene trees, resulting in a single basal polytomy and a significant amount of reticulation events across the whole extent of the phylogeny (Fig. S3). Pairwise hybridization numbers (*121*) also suggest a dominance of hybridization but are significantly biased by considering both micro and macroevolutionary processes.

The distribution of phylogenetic signal in phenotypic species measured using Pagel’s λ is also revealing (Figure S17). Although most likelihood ratio tests (LR) are significant, because specimens with similar phenotypes tend to aggregate in discrete lineages across the phylogenies (Supplementary_material), coherent groups are often interspersed by single specimens with discordant phenotypes. The most abundant and well-defined phenospecies have λ estimates consistently close to 1, whereas putative species that have intermediate phenotypes, and species that are clearly too broad (e.g. *P. circumalbata*) or some recently defined in phylogenetic terms (*P. micromarina, P. micromontana* and *P. microestepossa*) have lower λ values in at least one of the loci. The distribution of λ is likely influenced by imbalanced sampling size but serves to illustrate the extent to which intermediate phenotypes are also associated with intermediate genotypes.

Quantifying the contribution of hybridization and retention of ancestral polymorphism to incomplete lineage sorting (ILS) remains a methodological challenge in phylogenetics. Multispecies coalescent methods consider all ILS to predate species differentiation, while more complex models (*75*, *76*, *122*, *123*) are only usable when clear species concepts can be imposed *a priori*.

### Analyses of population structure identify ten partially admixed gene-pools which partially match observable phenospecies

An alternative to phylogenetic methods is to estimate population structure under mixture and admixture models, which can accommodate hybridization. To accommodate linkage information contained within loci we used the codon model of linkage implemented BAPS (*124*, *125*) in mixture and admixture models. The mixture analysis identified 10 evolutionary populations (gene pools, Figure 1b) consistently across starting conditions and the maximum number of populations. Mixture clusters show a high concordance with phenospecies and bGMYC clusters (figure 1a–c), although the concordance is, as always partial. The admixed fractions represent a large fraction of the phylogenetic signal (λ = 0.95–0.99), reflecting the similarity of different clustering results on the same data. The proportion of admixed individuals is quite low; 0.3 of individuals are assigned to a gene pool with a probability above 0.95, while in only 0.16 the largest fraction represents less than 0.9 of the genetic variation. The low admixture between gene pools suggests having strong post-mating isolation, as also does the high concordance between phenospecies and gene pools (0.77). The contribution of the different gene pools to the admixed signal is somewhat proportional to the overall representation of each gene pool (i.e. cluster size) in the dataset, but the linear relationship is not significant as clusters 3 and 4 have a larger contribution to the admixed signal than expected.

### The contribution to gene flow of gene pools is not uniform

The genetic connectivity between the ten gene pools was explicitly modelled using migrate-n (*126*) together with approximate demographic parameters (Figure 1d–e). The model is limited in that it cannot differentiate the different components contributing to lineage sorting, or better lack thereof. Because it uses a migration rate, it interpolates the transition between the imposed populations (gene-clusters) across genealogical time. The analyses were run using a Bayesian method and complete model. The estimated model parameters were mutationally scaled immigration rates (M) and effective population sizes (θ), average values have been reinterpreted in terms of migrants per generation, to filter out biologically relevant connections.

The contribution of the ten gene pools and their associated species to gene flow is not uniform, coherent with the general observation that hybridization propensity is not uniform across species and that patterns of hybridization are often dominated by a few species (*22*). Gene pool number five, strongly associated with *P. chalybaea*, is largely isolated from the rest, and only has a significant contribution to geneflow into pools six and four. The overall pattern of gene flow is strongly dominated by gene flow from to pools 1–4 associated with highly reduced endolithic species (Figure 1a,d–e). Clusters 6,7, 9 and 10 comprise a large fraction of species that are underrepresented in the dataset or that show intermediate morphologies and may be misrepresenting the underlying biological compartmentalization and are strongly interconnected between them. Cluster 8 which largely contains specimens of *P. variabilis* is the stronger contributor to geneflow into the reduced phenospecies, followed by cluster 7 which is loosely associated with *P. helygeoides* and *P. diphyodes* while cluster 9 is the one receiving most gene flow from them. In terms of mutationally scaled immigration rate, the stronger connectivity is found between pools 2 and 3 which are largely conspecific (*P. alociza*) and are clustered together in bGMYC (Figure 1), pool three is exclusively Iberian and could represent either resulting from population structure per se or be reflecting the assimilation of a preexisting species into *P. alociza* through introgression. Next to them the highest estimates of M is immigration into pools 6, 7, 8 and 9 from 4 and into 3,4,5, and 6 from 8.

The marked imbalance in gene flow to and from the reduced phenospecies and the abundance of intermediate morphs suggest that, despite the strong genetic signal, the evolution of Pyrenodesmia is strongly shaped by introgression and hybridization. While the group is likely formed by at least ten differentiated ancestral evolutionary populations or species, different genetic combinations contributed significantly to the observed genetic and phenotypic variability in the genus, making its systematic taxonomic treatment very difficult, and maybe even undesirable.

### Sequencing of mating type (MAT) idiomorphs identifies a large syngameon in European *Pyrenodesmia*

Despite the frequent claims of being integrative, “molecular taxonomic” surveys are often plagued by *post hoc* theorizing, because they use the same data for hypothesis generation and testing. To avoid this limitation we devised a method to identify reproductive compartments (i.e. biological species) by quantifying reproductive isolation at the dikaryotic stage, premating isolation in the sense of Steensels et al. (*90*). To do this we sequenced the MAT idiomorphs in samples containing dikaryotic mycelia and quantified their reciprocal association using: a) cophylogenetic methods, b) bipartite networks and c) unipartite networks. Each method provides a different perspective, but the unipartite networks at individual level provided the best framework to assign individual samples to reproductive units. We queried genome drafts of *Pyrenodesmia* for MAT cassettes and developed specific primers for two coding regions found in the complementary idiomorphs: the putative pheromone receptor of mat1-1 and the homeobox domain of mat1-2 (*127*). To simplify we will address them as alpha (α) and hmg respectively. The widely conserved, bipolar mating system and the widespread homothallism found in *Lecanoromycetes* greatly simplify the analyses.

We obtained sequences of either one of the idiomorphs in 0.88 of the samples; in 0.51 we obtained sequences of both mating types (Supplementary table 3), as in general we avoided including apothecia or apothecial primordia in the DNA extractions. The genetic diversity of the MAT idiomorphs in the reduced datasets is similar to those estimated for the phylogenetic loci (nucleotide diversity α = 0.06; hmg = 0.05), although haplotype diversity is higher (α = 0.681; hmg = 0.572) resulting in a high proportion of unique haplotypes (α = 0.66; hmg = 0.57).

The cophylogenetic signal in nucleotide datasets collapsed to haplotypes was evaluated using two closely related methods: RandomTaPas (*128*), which is recursive, and PaCo (*129*) which is a global method to interpret large tanglegrams. Both, identified low cophylogenetic signals with normalized Gini coefficients (G*) of 0.76, which, interpreted as a fraction of dissimilarity, is roughly the inverse of the 0.15 concordant fraction found between observed and randomized phylogenies for the phylogenetic loci. Cophylogenetic methods summarizing multiple random tanglegrams contain very few well-supported cophylogenetic relationships, suggesting the absence of a coevolutionary association between both mating type idiomorphs. This can also be caused by the alteration of genetic drift due to the suppressed recombination around the mating type cassette (*130*).

To better model reproductive isolation, we made use of network models implemented in R packages *bipartite* (*131*) and *igraph* (*131*, *132*). Under the premise that mate recognition and compatibility function at protein level, we translated the nucleotide sequences and collapsed them to 0.99 similarity haplotypes (*133*). Additional analyses using nucleotide and aminoacid haplotypes were also carried out. Because they differ in their resolution but not in the overall pattern and are left out of the manuscript.

Bipartite networks use the haplotypes of the alpha and hmg idiomorphs as independent strata, giving two disjoint sets of nodes. The edges quantify the times each combination of haplotypes is found in the dataset. In unipartite networks nodes represent specimens and edges quantify the number of aminoacid haplotypes of any of the two idiomorphs number shared between specimens.

All estimated networks show a similar pattern of compartmentalization, with a single subnetwork grouping most of the nodes, that is 0.69 in the bipartite network of haplotypes, 0.87 in the bipartite network at 0.99 similarity and 0.89 in the unipartite network (Figure 1f). The rest of the subnetworks group underrepresented sequences, species and geographic regions. The number of compartments is proportional to the similarity threshold used to compress the dataset, decreasing from 66 to 22 when collapsing the aminoacid sequences to 0.99 similarity (Figure 2, Supplementary material). The statistical properties of the complete network, obviously arise from those of the main interconnected subnetwork. Both have very low density (0.036, 0.045), low average centrality (0.059, 0.074) and an average path length of (3.654, 3.656) (Supplementary table X). The distribution of statistical properties at the network level is not homogeneous between species, and nodes identified as *P. alociza* and *P. erodens* tend to have a higher centrality and degree than the rest, while betweenness is similar across the network.

**Fig. 2.**
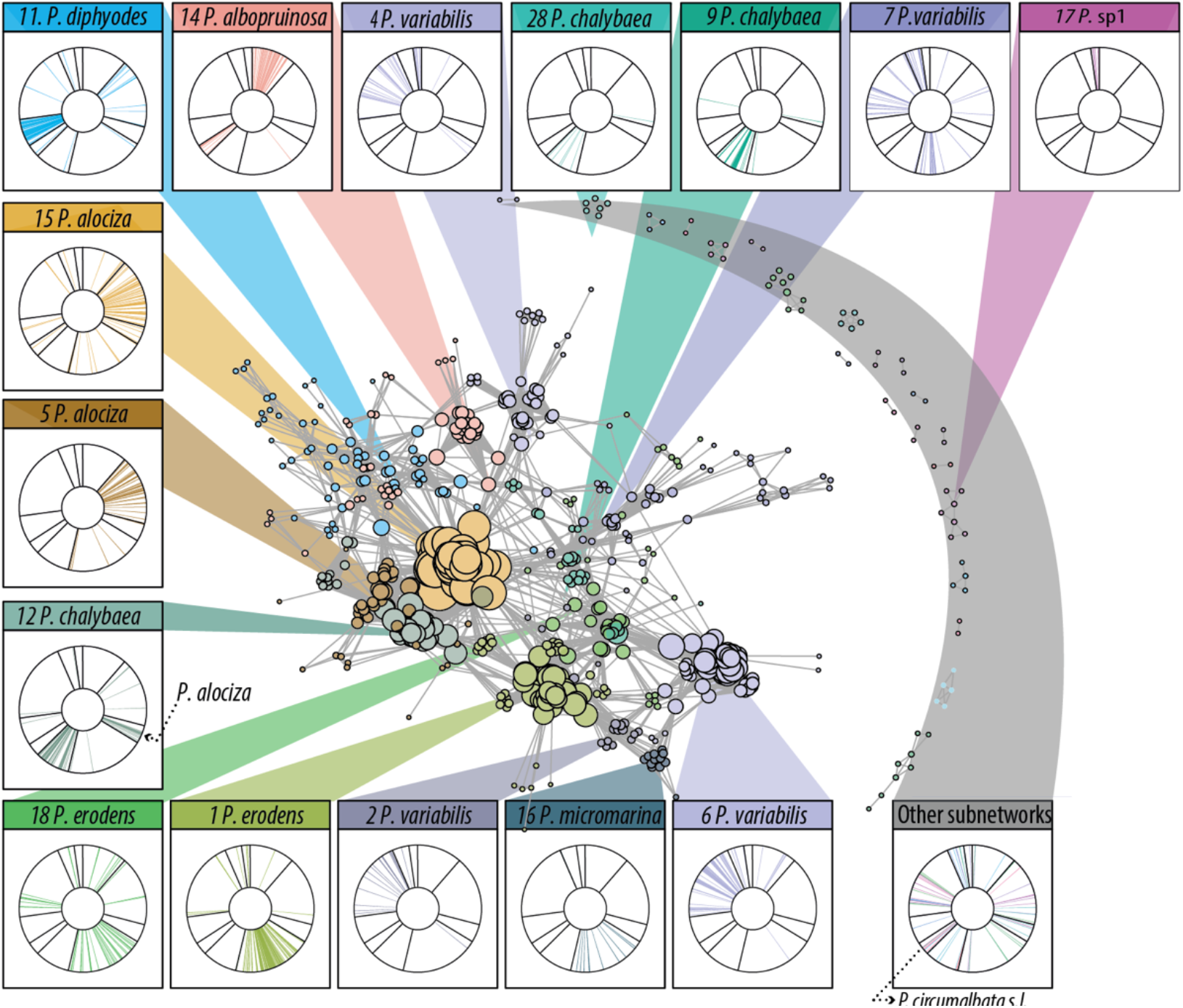
Modularity of the unipartite network of MAT idiomorph cooccurrence. Figures a-c and f provide a graphical summary of the structure of the dataset using the assignment of specimens to gene-pools estimated in BAPS as organizing scaffold. estimates provided in the supplement. The network was simplified to include only strong connections (2Ne>2). **f**| Assignment to Mating compartments using the unipartite network of MAT idiomorph cooccurrence. The network contains a main subnetwork comprising most specimens and 20 secondary ones.

The presence of well-supported reproductive units (modules) was further investigated using the Louvain (*134*) community detection algorithm on the unipartite network. The performance of different algorithms was evaluated using Bayes Factors and two null models using R package *Robin* (*135*). The selected Louvain algorithm estimated a modularity of 0.77 for the complete network, and split the main compartment into 13 modules, adding up to a total of 34 modules (Figure 2), which group between 0.11 and 0.05 of the specimens. Their assignment to modules is quite coherent with the results of population genetics and bGMYC, but the overall association estimated using Cramer’s V between phenospecies and modules (0.47), and between modules and phenospecies per geographic region (0.57), are weaker than that reported between phenospecies and gene-pools (0.77), despite having a larger number of categories.

The bipartite networks are less useful to assign specimens to single reproductive compartments but allow addressing whether asymmetries in connectivity between both strata. The 0.99 bipartite network is very similar to the unipartite network in terms of compartmentalization (21 compartments). In both cases, the diversity of the alpha locus (Supplementary table 3) is slightly higher than that of hmg, which results in a slightly asymmetric network. (−0.19). The c-scores show disaggregation in both levels, but the checkerboard analyses are biased by the abundance of zero values. The V.ratios give a better representation of aggregation and show that the hmg stratum is more aggregated (6.18) than alpha (1.39), which is slightly higher than the numerical imbalance between the number of haplotypes in both strata (hmg/ α = 91/136 = 0.67). The higher mean number of shared partners (hmg/ α = 0.07/0.11 = 0.64) and partner diversity (hmg/ α = 0.9/1.45 = 0.62) found for the hmg locus likely reflect this numerical imbalance. Contrarily, the mean number of links per sequence is much higher in hmg (8.8) than in alpha (3.93) and differs (0.45) from the reported asymmetry expected as a result of haplotype richness. This asymmetry is likely to result from a net difference in specificity between both strata, as reflected by hmg having a much higher number of realized links than alpha, as shown by the net difference in cluster coefficient (hmg = 0.065, α = 0.043), although when weighted using Ophal’s method based on four-paths, cluster coefficients are higher in alpha (0.20) than in hmg (0.12) due to the imbalance between both strata.

In summary, the study of mat loci show that *Pyrenodesmia* forms a complex syngameon in Europe in which species keep their reproductive boundaries open and are able to hybridize with each other. Mating happens randomly across the dataset, but at the same time is highly structured. It is not reasonable to consider mating as equally probable across all specimens in the dataset, because geographic and ecological boundaries, local vicinity and population sizes contribute significantly to premating isolation and have not been explicitly accounted for. The randomness in mating is supported by the lack of cophylogenetic signal found at nucleotide level, and most importantly by the lack of significant difference between the observed unipartite network and the most conservative null model, a random graph (dk-series 2.1 model), which preserves the joint degree distribution and the clustering coefficient of the original network but not the full clustering spectrum (*136*). Overall, there is a clear preference for intraspecific mating, or mating within gene pools (Supplementary_material), which is obviously limited by the local occurrence of both species and genotypes. Furthermore, the discordance found between the promiscuity identified in dikaryotic samples and the gene-pool structure suggests that premating reproductive barriers are weaker than post-mating barriers in *Pyrenodesmia*. A thread of doubt does remain, because it is possible that the post-mating hybridization identified in the sanger dataset reflects the presence of non-viable dikaryotic combinations caused by the lack of mating isolation, and not actual recombined hybrids.

### Phylogenomic reconstructions identify signatures of both hybridization and hybrid speciation

To address this discordance, we looked for signals of hybridization at a genomic scale. We assembled a comparative genomic dataset including 23 specimens of *Pyrenodesmia* and two external references (Supplementary_material). Their phylogenomic examination was carried out using 2767 single copy orthologous genes, whose phylogenetic trajectories were modelled independently using maximum likelihood (ML) and bootstrapped replicates for nodal support. The summary Figure 3d combines the topology of a cluster consensus network (Figure S4) and the nodal support values of a consensus phylogenetic tree (Figure S5). Both consensus approaches allow identifying a phylogenetic signal of hybridization at whole genome level. The estimation of pairwise hybridization networks (*121*) identifies 6 hybridization events across the ML reconstructions, which is coherent with the reticulations depicted in the cluster consensus network. Three of them are intraspecific, while two link specimens of *P. alociza* with non-endolithic species, some ill-defined reduced morphs. A third interspecific reticulation is basal to a wider clade containing *P. erodens*, which is otherwise well supported with 0.92 internode and tree certainty values.

**Fig. 3.**
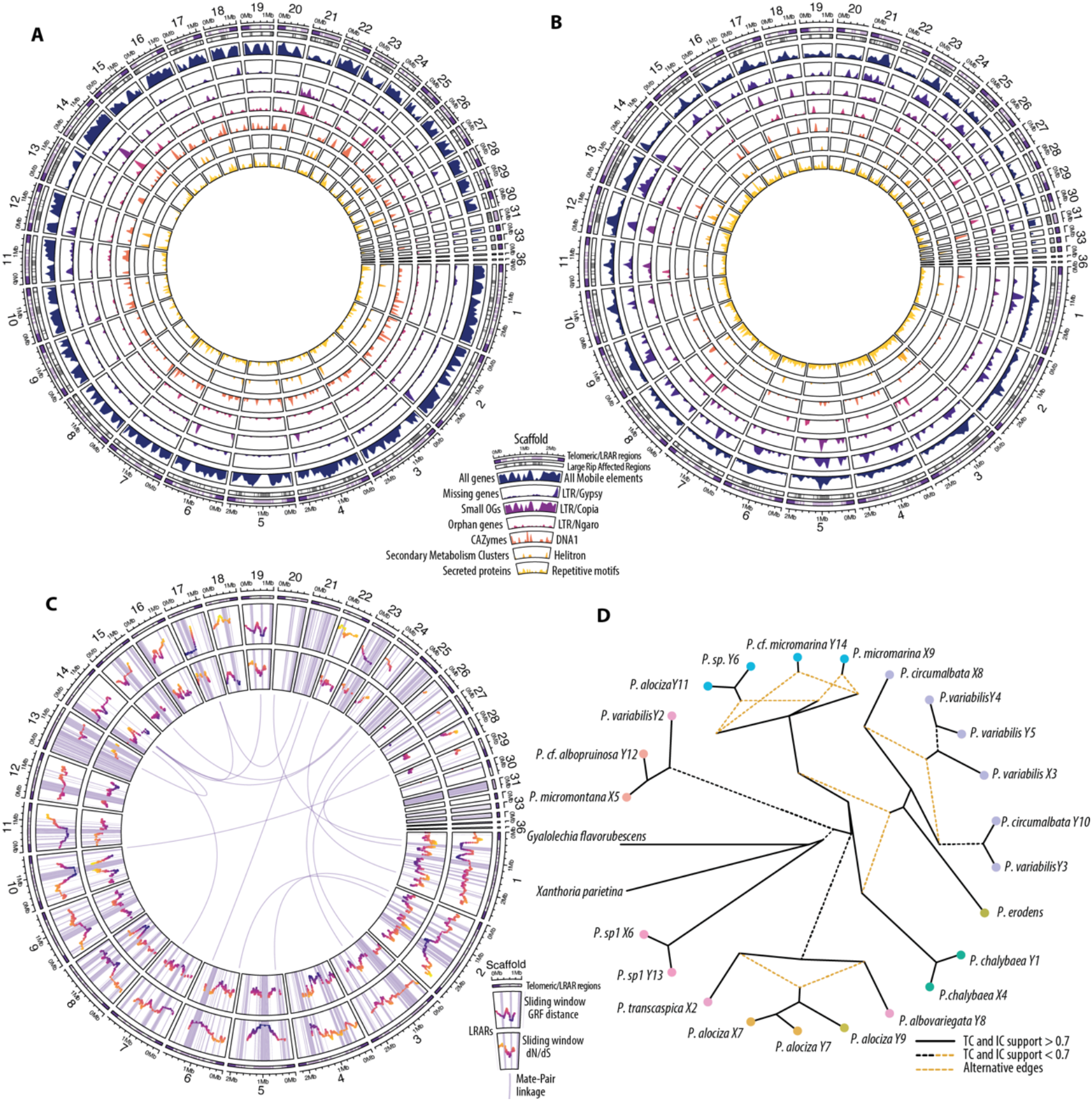
Summary of the genomic draft of *Pyrenodesmia* erodens and its phylogenomic context. **a**| Genome structure and annotation. The plot represents the individual scaffolds numbered from 1 to 36 and their length. Each sector contains the following fields: 1| Regionalization. Telomeric and subtelomeric regions of 0.35Mb from the terminus are highlighted in darker color, large RIP affected regions (LRARs) and 0.03Mb flanks are highlighted in a lighter color. 2| LRARs identified using theRipper. 3| Density plot of coding regions calculated using a sliding window of 50Kb, common to all other density plots. 4| Density of missing genes, it considers the syntenic position of genes missing in the syntenic blocks of *P. erodens* but present in more than X other Pyrenodesmia genomes. 5| Density of genes assigned to small OGs) found in at least 5 genomes. 6| Density of singleton or orphan genes, assigned to OGs found only in *P. erodens*. 7| Density of genes related to Carbohydrate metabolism (CAZymes) 8| Density of Secondary metabolism clusters (Terpenes, PKS and NRPKS) 9| Density of secreted proteins. **b**| Annotation of repetitive and transposable elements in the *P. erodens* genome. Each sector contains nine fields, the first two are the same as above, the rest show the estimated densities of: 3| All mobile elements. 4| LTR/Gypsy. 5| LTR/Copia. 6| LTR/Ngaro. 7| DNA elements including those of Voyager proteins 8| helitron 9| repetitive sequences. **c**| Summary of the phylogenetic landscape across the *P. erodens* genome. Each sector representing a scaffold contains three fields: 1| regionalization of the genome. 2| Generalized Robinson-Foulds distances between neighboring single copy orthologous genes (SCOGs). Each data point represents the average distance between the ML phylogenetic reconstructions on overlapping sliding windows containing 20 consecutive SCOGs. Vertical bands highlight the identified LRARs including centromeric regions. 3| Sliding window of dN/dS values calculated using the same method as above. On both the numerical values are emphasized by using a color gradient between blue and orange. The central region of the plot shows the linkage between scaffolds using regions with a high density of split mate-pairs, in which each paired sequence aligns to a different scaffold. **d**| Phylogenomic consensus network. Well supported edges (IC and TC > 0.7) are drawn as whole black lines, edges common between the consensus tree and network reconstructions but with lower support are shown as black dotted lines. Edges representing a hybridization event, with IC/TC values are highlighted as colored dotted lines. The color of each tip is coherent with the assignment to the gene pools in Fig1b.

### *P. erodens* shows signs of genome concertation and stabilization

The evolutionary importance of hybridization has been recently reconsidered in whole genome surveys, which found evidence of cross-specific gene transfer across the eukaryotic tree of life, contradicting the canonical view that hybridization is a rare and deleterious event. Hybridization in fungi (*90*), originating through sexual or parasexual gene transfer (*137*), has been identified as a major cause of genomic instability (*138*) but has also been proposed as an important source of adaptative traits and genetic variability (*20*). What is still not clearly understood is the outcome that hybridization has in haploid organisms, in which the vegetative outcome is not necessarily a one-to-one hybrid, but a complex gradient of recombined genotypes. Additionally, chromosomal structure may determine a mosaic of regions with enhanced and suppressed recombination, and genome defence mechanisms may determine asymmetries in the stability of gene content in hybrids.

We addressed this heterogeneity using the genome of *P. erodens* as baseline reference. Despite the strong contribution of this species to the overall admixed signal, and mating network, the phylogenomic reconstruction suggest that its genome has been shaped by an old hybridization event. Its evolutionary placement is quite coherent across loci and seems dominated by mechanisms ensuring the stability of its gene content. With 41.8Kb its genome is 0.3 larger than the quasi-complete reference genomes of *X.parietina* (14) and *G. flavorubescens*, but contains roughly twice the number of chromosomes (24,14 and 11.5 respectively) approximated counting telomeric (CCCTAA/TTAGGG)n motifs, 14 on both ends, 20 on one end in *P. erodens*. The genome size is not significantly different to other Pyrenodesmia draft genomes, although their sizes may be smaller due to the stringent metagenome filtering used.

The difference in chromosome number and the similarity in genome sizes suggests that genome duplication or aneuploidy may have shaped the evolution of the whole genus. More contiguous assemblies would be needed to thoroughly discuss the extent to which this is shared among *Pyrenodesmia* species or reflect older events taking place within the subfamily *Caloplacoideae*. The ancestral origin of the hybridization event is also supported by the low of heterozygosity found and the low levels of interchromosomal homology observed, with a single intragenomic syntenic region between chromosomes 15 and 24, related to mobile elements, and the lack of shared k-mers between scaffolds.

Despite being similar in size, the genome of *P. erodens* has a lower gene content than the other high contiguity genomes in the dataset (Supplementary_material). This slight depauperation is also observable in the functional annotations, where the tendency towards a reduction of genes identified per functional unit (e.g. pfams, interpro, COGS, etc.) is clearly observable (Figure 4b) although it is partly obscured by the overall stability in gene content (Figure 4a). The simplification of the genome also results in a reduction on the number of tRNAs (46) compared to the references (57), which could also explain the loss of genes with discordant codon usage. The rest of *Pyrenodesmia* genomes show similarly reduced tRNA counts, except for *P. alociza* which has consistently higher tRNA counts (77–137), correlated with the strong signal of hybridization found for all its specimens in the phylogenomic dataset.

**Fig. 4.**
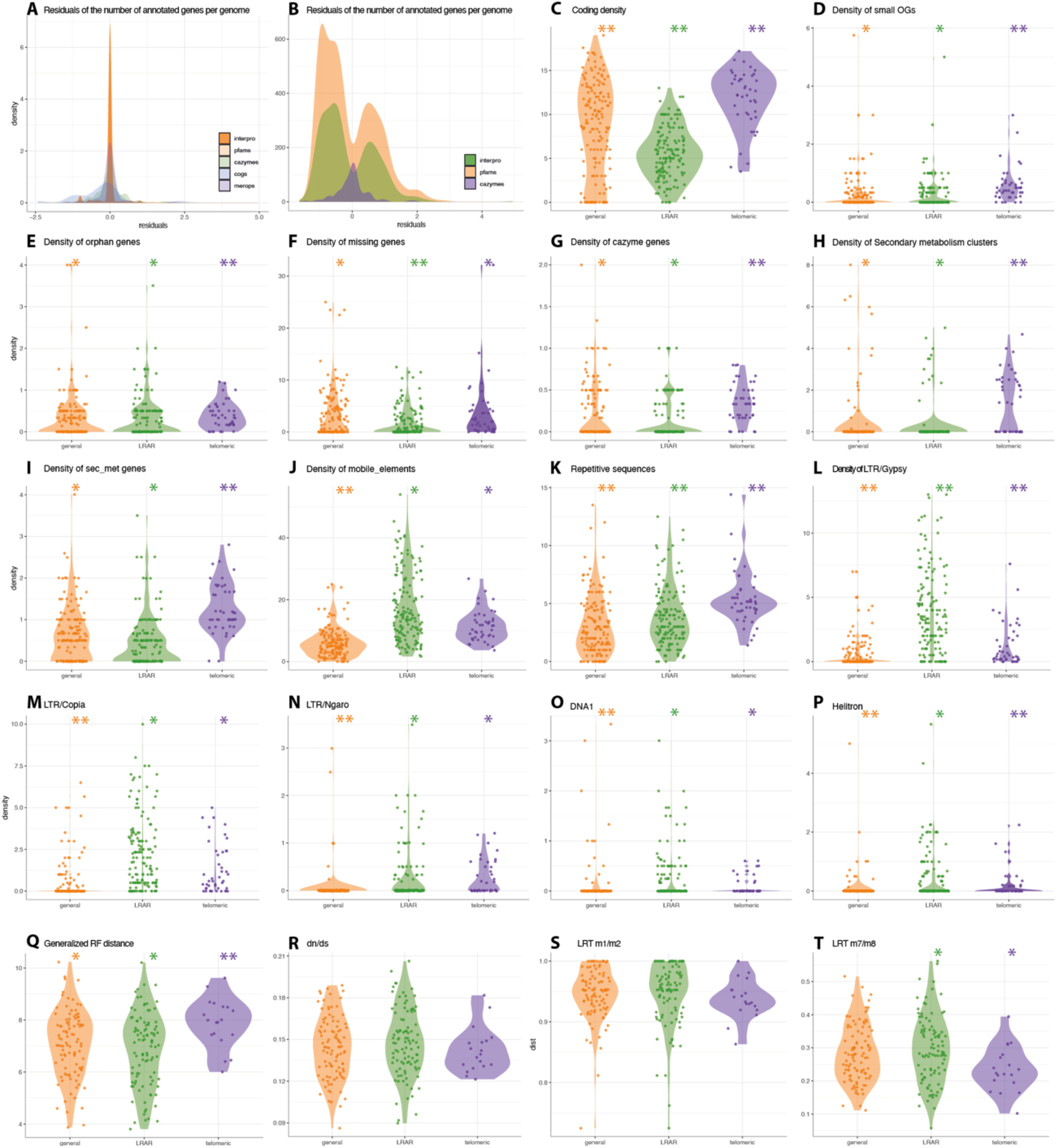
Intragenomic regionalization of the *P.erodens* genome. **a**| Density plot of the standardized residuals of annotated gene representation in the genome. It includes all genes which rendered a positive match to Pfam, Interpro, COG, Merops and DbCAN databases. The expected value was imposed as the median of all other Pyrenodesmia genomes. **b**| Density plot of non-zero standardized residuals of annotated genes. Highlighting the towards depauperation of gene content. **c-p**| Show the density of different genomic features in the three imposed regions, telomeric and subtelomeric, LRARs and their flanks and the rest of the genome. Densities were calculated using a sliding window of 20Kb: **c**| Coding density. **d**| Small OGs. **e**| Orphan genes (singleton OGs). **f**| Missing genes. **g**| CAZymes. **h**| Secondary metabolism clusters. **i**| Secreted proteins. **j**| mobile elements. **k**| Repetitive sequences **l**| LTR/Gypsy. **m**| LTR/Copia **n**| LTR/Ngaro **o**| DNA1. **p**| Helitron. **q-t**|Distribution of phylogeny based metrics averaged across groups of 20 neighboring OGs. **q**| Generalizaed Robinson-Foulds distances. **r**| Ratio of non-synonymous and synonymous substitutions (ω=dN/dS). **s**| Averaged value of the Likelihood Ratio Test between models m1, which considers loci being either conserved or neutral (ω<1 & ω=1), and m2, which include a third category allowing loci to be under positive selection (ω<1, ω=1 & ω>1). **t**| Averaged value of the Likelihood Ratio Test between models m7 and m8. Both models consider that ω varies across sites following a beta distribution restricted to the interval (0,1), model m8 includes an additional category of sites under selection which can take a value of ω above 1.

### Genome defence mechanisms are largely responsible for the simplification of the *P.erodens* genome

The dataset shows significant conservation of syntenic blocks, but loss of individual genes is widely distributed across the genome. Most notably, the genome of *P. erodens* is largely affected by Repeat Induced Point Mutations (RIP), a genome defence mechanism largely responsible for preventing the proliferation of transposable elements (TEs) in fungal genomes. Because RIP is a widespread mechanism in fungi, the GC content (0.44) and the proportion of the *P. erodens* genome affected by RIP (0.28) are within the values observed in other Ascomycota (Supplementary_material). What makes this genome peculiar that most RIP is found within large RIP affected regions (LRARs), adding up to 0.27 of the total genome size. Also, these LRARs are unevenly distributed among the scaffolds (Figure 3a–c), eight of which (11–13, 15, 17, 21, 23 and 25) are disproportionately affected by RIP (Figure 3a–c) to the extent of having very low coding density (Figure 3a), a high density of genes missing from syntenic blocks (Figure 3a), identified as single-copy orthologs (Figure 3c) and hardly any gene with a functional annotation (Figure 3a). These “supernumerary scaffolds” do not simply represent assembly artifacts, they are long (1.54–0.7 Mb) and amount to 0.22 of the total genome size.

The canonical view is that LRARs result from the proliferation of TEs and the subsequent genome defence. In addition to that, we hypothesize that RIP is a strong force driving genome stabilization and concertation, if not by itself, coupled with other mechanisms of genome defence and RNA interference. This idea is coherent with the evidence that fungal polyploids are most obvious in yeasts defective in RIP associated genes (*139*). RIP eliminates non-contiguous homologous chromatin regions which are repeated. In its nature RIP is symmetric, resulting in the deletion of both copies in a heterokaryon. What we observe, however is the maintenance of genes that largely share a common phylogenetic history, a directionality that could be explained by a hypothetical interaction between RNA interference mechanisms and RIP. Beyond hypothetical considerations, this directionality can be observed in the strong association between LRAR and mobile element density in the described “supernumerary scaffolds”. The causal relationship is however difficult to stablish because most of the signal is caused by the presence of LTR/Copia and LTR/gypsy elements which attach specifically to GC poor regions such as centromeres. It has been hypothesized that the proliferation of repetitive elements is also promoted by hybridization, but the extent to which LRARs originate from LTR-mediated extension of RIP affected regions or are the original cause of RIP is difficult to assess.

Moreover, LTR elements are typically found in centromeres, and although we attempted classifying LRARs according to their content in transposable elements, we did not manage to properly differentiate centromeres from secondary LRARs. Finally, RIP tends to extend beyond the homologous region potentially altering flanking genes. In *P. erodens* we found a strong collinearity between the density of genes missing from syntenic blocks and the density of transposable elements using sliding windows (Supplementary_material) as well as a negative correlation with the total gene density. This correlation is stronger when telomeric regions are excluded, but is not statistically supported when windows with 0 genes are taken out. Overall suggesting a strong association between the loss of genes and the action of RIP on the genome.

### RIP also drives genome compartmentalization and evolutionary stratification

While mobile elements and genome defence mechanisms play a major role shaping the gene content of *P. erodens*, they are also a major source of evolutionary stratification. We surveyed evolutionary stratification by mapping the discordance of phylogenetic signal between contiguous single copy OGs using generalized Robinson-Foulds distances on overlapping sliding windows (Figure 3c). This approach shows that regions proximal to putative centromeres, especially those placed between two large LRARs show lower average phylogenetic discordance between OGs (Figure 3c), as well as an accumulation of non-synonymous mutations, shown as higher average dN/dS values (Figure 3c), and divergent average m1/m2 and m7/m8 ratios (Supplementary_material). This is coherent with the idea that areas proximal to centromeres show suppressed recombination (*140*–*143*) and significantly contribute to genome architecture (*144*). The proliferation of LRAR caused by the interaction of genome defence mechanisms (RIP) and the proliferation of mobile elements in AT rich regions, results in the development of polycentric chromosomes (*145*), which behave locally in a similar way as reported for more complex repeat based holocentric (*146*) chromosomes in plants. The proliferation of transposable elements is also increased in subtelomeric regions, which are also affected by RIP but show significant differences in behaviour and gene content compared to the rest of the genome.

To study the differences across genomic regions we split the genome into three functional regions: subtelomeric, LRAR and general. The first category includes 350Kb proximal to scaffold ends containing telomeric, including LRARs found within them. The second contains centromeric and non-centromeric large RIP affected regions including 30Kb on each flank. The third contains the rest of the euchromatin. These three regions differ significantly in their evolutionary and functional stratification (Figure 4). The subtelomeric regions have slightly higher average RF distances between single copy orthologs (Figure 4q), and although the dN/dS values (Figure 4r) are also lower the difference is not statistically significant. Meanwhile, while the difference between LRARs and the rest of the genome is not significant, the LRARs show a more bimodal distribution of metrics reflecting the presence of LRARs centromeric or proximal to centromeres and other regions that do not show a strong signal of reduced recombination, because they are younger or incorporate genes that relocated within TEs.

The three regions differ in the extent to which they have been affected by different processes. The coding density is obviously lower in LRAR flanks than in the rest of the genome, while subtelomeric regions although often including LRARs have a slightly higher density than the rest (Figure 4c). Rare genes belonging to small OGs and orphan genes are significantly associated with subtelomeric regions (Figure 4d–e), while the density of missing genes is obviously higher in LRARs (Figure 4f). In contrast, while the three regions differ in the overall density of repetitive sequences (Figure 4k), the density of mobile elements is statistically similar in subtelomeric and LRAR regions (Figure 4j), a divergence that may result from the different extent to which the three regions are affected by the proliferation of different types of TEs (Figure 4l-p).

Finally, the telomeric regions have a significantly higher density of genes related to carbohydrate metabolism (CAZymes, Figure 4g), secreted proteins (Figure 4h), and secondary metabolism clusters (Figure 4i) than the rest of the genome. Phylogenetic stratification of the genome is also observable across the dataset. Regions of reduced and increased recombination are intermixed across the genome (Figure 3c) and are strongly associated with the presence of LRARs. This suggests that subtelomeric regions, which have increased recombination and are strongly associated with certain mobile elements, are key in the acquisition of functional diversity and adaptive traits. Furthermore, subtelomeric regions show higher average dissimilarity between single-copy OG phylogenies than the rest of the genome (Figure 4q) and slightly lower evidence for positive selection (Figure 4r–t). The scale at which stratification takes place is finer than the coarse regions used in this survey and its study requires an in-depth systematization to characterize the recombination history of individual chromosome regions.

## 3. DISCUSSION

Language provides us with the basic tools to imagine, organize and share our knowledge of the natural world. Taxonomists use a hierarchic system of linguistic categories to represent and organize the observable discontinuity of organic variation. For them, life is organized in discrete units, species, that are subject to a continuous process of evolutionary change and are connected through their shared ancestry.

Reconciling the continuous nature of evolutionary change with the discrete outcome of speciation is a foundational challenge in evolutionary biology, which is becoming ever more apparent as the evidence that cross-specific exchange of genetic information is frequent piles up across all domains of life. Viewing species as only units of biological organization is often a useful compromise; flexible enough to allow interpreting lack of reproductive isolation in terms of interspecific hybridization and introgression. But to what extent is it a harmful simplification? Is it necessary to perpetuate an unrealistic *scala naturae* made up of fix categories and deterministic expectations? Life is a complex evolving system in which both genealogical and non-genealogical bonds are equally relevant in shaping its structure, diversity, variability, and stability.

The lichen genus *Pyrenodesmia* provides a clear example of a genus whose species remain interlinked at a supra-specific level. The genus contains a discrete number of identifiable genetic entities which associate significantly with already described morphospecies, as well as several morphologically and genetically discordant specimens which amount to ca. 30% of the dataset. Some of them are unrecognized species, but others are clearly intermediates in their phenotype, their genotype, or both. The evidence that the identifiable genetic groups transfer genetic information with each other, as provided by admixture and gene flow (migration) analyses (Figure 1a–e) could be interpreted in terms of hybridization or introgression between species resulting from secondary contact between well-differentiated species. However, in the European continent, all species are interconnected through mating in a single large reproductive network, a syngameon, despite which the main species remain largely distinct and distinguishable. Moreover, the phylogenetic (Supplementary_material) and phylogenomic trees (Figure 3d, Supplementary_material) reflect the presence of a long-standing history of genetic exchange more than a series of unresolved secondary contacts.

Coherently with findings made on other syngamea, the contribution of the different species and gene pools to the hybrid signal is asymmetric at pre- and post-mating levels. In Pyrenodesmia, cross-specific bonds seem to be driven by species that have reduced endolithic thalli (Figure 1a–f, Figure 2), and appear to be of hybrid origin in the phylogenomic reconstructions (Figure 3d), particularly *P. erodens* and *P. alociza* (gene pools 2,3). These are preferentially introgressed by larger epilithic species of gene pools 6–9). The observed asymmetry es even stronger when considering migration between gene pools using each mat idiomorph. The directionality of geneflow as estimated using migrate-n is difficult to interpret, especially because there is some discordance between gene pools and phenotype groups. Further evidence of the asymmetry is provided by the net difference in average closeness centrality estimated for the different mat modules (Supplementary_material), and the strong contribution of *P. alociza* and *P. erodens* modules to the network structure.

In terms of evolutionary ecology, there is no clearer contributor to fitness than mating itself. In that respect, syngamea could be interpreted to behave in a way analogous to metapopulations (metaspecies), where asymmetries in mating are strong drivers of both population structure and local adaptation. Contrary to metapopulations, geneflow between species may result not just in the acquisition of maladaptive traits, but have stronger selective consequences. In that respect, the diversity of reproductive interactions can be interpreted in terms of reproductive niche, transposing an Eltonian niche concept (*147*) to describe the width of the genealogical vicinity with which mating is possible. Seen from that perspective, mating behaves the same as any other biotic interaction. Species will show different degrees of reproductive specificity, from a very narrow reproductive niche as *P. chalybaea* to a generalist with fully open reproductive boundaries. An idea subsequent to that of niche, which is sometimes strongly discussed by ecologists, but is a central consideration is that the observed interactions result from the realization of a broader fundamental niche which may result from local processes such as competition or density. This would reconcile the discordance between the lack of compartmentalization and the strong association between the estimated modules, phenotypic species and region.

The outcome of reproductive interactions may also be interpreted in terms of their effect on both mates, which can be beneficial by increasing the transfer of adaptive traits to deleterious in case of, for instance genome or mitonuclear incompatibilities. In that respect we hypothesize that the strategy of the endolithic *Pyrenodesmia* species is analogous to sexual parasitism (*148*), which use closely related species to propagate, at the expense of certain gene leakage between species and some backcrossing leading to hybrid phenotypes. It is not necessary to annihilate the accepting genotype to be parasitism, but simply to increase its own fitness at the expense of the other species in the syngameon. Compared to other models of sexual parasitism, fungi are different because of the meaning of mating and the haploid nature of the vegetative mycelium. The outcome of hybridization is not completely known, but it is not necessary Mendelian, especially because hybrids result in a whole probabilistic spectrum of partially recombined haploid genotypes.

As genomes diverge and species within the syngameon diversify, hybridization comes at a stronger cost to both sides of the interaction. The genomes of sexual parasites must reflect the deleterious effects of non-specific mating at genome level but also the traits selected to maintain the feedback loop in which maintaining the increased fitness of the parasitic behaviour overweighs any other adaptive strategy.

In the phylogenomic dataset we observe two species which show different traits related to their hybrid history. The genome of the *P. erodens* culture seems to reflect the long term outcome of hybridization. Its genome shows traces of genome stabilization. We do not know whether the higher chromosome number results of hybridization or is a shared trait within the genus. Its gene content remains quite stable despite hybridization, although partial gene loss is observable across the whole genome. The loss of genes is mostly caused by the action of genome defence mechanisms, especially RIP, which disproportionately affects certain scaffolds and chromosomes to the point of completely obliterating their gene content. This asymmetry is not typical of RIP and cannot be explained by the observed proliferation of transposable elements alone. This makes us conclude that RIP, in interaction with other genome defence mechanisms is a key factor maintaining fungal genomes haploid and driving concertation of gene content in hybrid genomes.

The accumulation of large RIP affected regions (LRARs) also modifies the recombination landscape of the *Pyrenodesmia* genomes, because it generates regions of reduced recombination between LRARS that likely behave as centromeres (Figure 3c). These provide certain protection from the deleterious effects of recombination and may contribute to the overall stability of the genome but are on the long run prone to accumulate non-synonymous mutations (Figure 3c). Meanwhile the subtelomeric regions of the genome remain active both in terms of recombination and incorporation of mobile elements, retaining a strong potential for the acquisition of functional variability (Figure 3–4). The loss of certain genetic elements like heterokaryon incompatibility genes, as observed in *P. erodens* is a potential mechanism to increase cross-reproductive fitness, while the reduction in tRNA content may provide additional transcriptional silencing of foreign genes.

The genomes of *P. alociza* on the other hand appear as recent hybrids. They show increased genome content and tRNA count, but are difficult to interpret because they do not originate from a haploid tissue and may not represent accurately the real genomic structure. They do hint to a first stage in hybridization in which genome stabilization is still an ongoing process, which requires a more in-depth look.

On the whole, this survey raises worrying questions on the way phylogenetic surveys interpret the genetic variability of fungi. Is the supraspecific evolutionary unit found in *Pyrenodesmia* an exception? Most likely it is not, and while molecular characters provide an objective description of the relatedness between genes, and organisms, the methods used impose very strong opinions on the interpretation of relatedness, the outcome of evolution and the properties of populations and species. To what extent are we doing a good job describing fungal evolution in terms of hyperdiversity and cryptic speciation? How much are we affected by confirmation bias? Or are we assuming simplistic operational expectations because they are publishable?

## 4. MATERIAL AND METHODS

### Dataset assembly, collections and morphological identification

We assembled a collection of *Pyrenodesmia* specimens growing mainly on limestone surfaces across the Mediterranean region of the Eurasian continent. These were enriched with reference collections made in Central Asia and North America. Sample location is shown in ST 1. The identification of the specimens using morphological characters is a daunting task. The univocal identification of specimens to one taxon is often hampered by the lack of clear differentiation, the abundance of intermediate forms and the difficulty to carry barcoding using molecular characters. In general terms we used the monographic work of Wunder (*96*) as a backbone over which we integrated newer taxa and taxonomic concepts (*97*, *99*, *101*, *102*, *105*). This effort has been largely summarized by Frolov et al. (*106*).

### Development of specific primers

The assembly of a multilocus dataset proved particularly challenging. Added to the species identification problems, the microscopic nature of most species hampered the process of DNA extraction. This is especially true for endolithic samples which have low fungal biomass and high amounts of Calcium carbonate. In most cases the yield sufficed to amplify repeated loci as ITS or mtLSU, but not single-copy nuclear loci, a problem exacerbated by the mediocre performance of standard fungal-specific primers. To enable de assembly of a multilocus dataset we developed specific primers for *Pyrenodesmia*. Instead of developing primers internal to those used as standard, we aimed at providing Teloschistaceae/Pyrenodesmia-specific primers covering a larger genomic region, more in tune with modern PCR and sanger sequencing capabilities. For this we assembled nine reference genomic drafts to serve as reference and to test the usability of genome-wide phylogenomic surveys. We queried the assembled loci using Blastn (*149*) as to extract complete sequences of a set of loci commonly used in previous phylogenetic surveys at generic and family levels: The mitochondrial large subunit rRNA (mtLSU), the nuclear Internal transcribed spacer region of the rRNA cistron (ITS), and five protein coding loci: mini-chromosome maintenance complex protein 7 (mcm7), Elongation Factor α (Efα), β-Tubulin, and the largest subunits of the RNA polymerase II (RPB1 and RPB2).

Further subsetting, multiple sequence alignment and primer design were carried out in Geneious v7 (*150*) using the included augustus (*151*) and Primer3 plugins. Multiple sequence alignments contained a single gene copy per genome with the exception of ITS and mtLSU, and all belonged to contigs predicted as fungal. To provide more specific primers for mtLSU, which is prone to cross-contamination, and to exclude the photobiont sequences in ITS we included contaminant sequences of lichen-associated fungi and *Trebouxia* photobionts as found in the sequenced metagenomes.

### PCR and sequencing

Because standard PCR protocols did not deal well with the low quantities of DNA used, PCRs were carried out using the KAPA3G Plant PCR Kit (Kapa Biosystems, VWR) using the following cycling conditions for the phylogenetically informative loci: Initial Denaturation at 95°C for 5 minutes, 40 cycles of Denaturation at 95°C for 20s, Annealing at 57°C for 15s, Extension at 72°C for 30s and a Final Extension step at 72°C for 7m. For the MAT loci 40 cycles of Denaturation at 95°C for 30s and annealing at 60°C for 15s were used instead.

All reactions were carried out using 10 μL reaction volumes. The mastermix was modified to include: 5 μl 2x KAPA Plant PCR Buffer, 0.7 μl Forward Primer (10 μM) and 0.7 μl Reverse Primer (10 μM), 1-3μl Template DNA, 0.05 μL KAPA3G Plant DNA Polymerase (2.5 U/μL), fill up to 10 μl with PCR-grade water. Sanger sequencing was carried out by Microsynth Austria on an ABI 3001 platform.

### Dataset assembly

Although more than 1200 specimens were processed, the final dataset includes 824 specimens for which we could assemble a credible and almost complete data-matrix (98%), with sequences of at least three out of the five loci used. To achieve the maximum completion of the dataset we repeated the amplification and sequencing progressively increasing the amount of template until the gaps were filled or the extract was exhausted. The microscopic nature of most samples did not allow doing multiple extractions per sample; in consequence sequences are never pooled from parallel extractions. The dataset used in the manuscript does not make use of mtLSU nor RPB2 sequences, but we provide for coherence with previous surveys that may have RPB2 or mtLSU. The mitochondrial LSU was discarded as it contains two complementary homopolymeric regions that degrade the performance of Sanger sequencing beyond the limits of usability. Additionally, we suspect the complex reproductive biology to cause significant mitonuclear discordance and heteroplasmy. The two overlapping fragments of a long region of RPB2 were the last to be sequenced and remain very incomplete due to the exhaustion of DNA extracts.

All sequences were processed in Geneious v.11 (*152*) departing from the raw .ab1 files. After a first alignment using mafft (*151*), sequences were trimmed, and base calling errors were manually corrected. Correction was aided by the simultaneous graphic visualization of the electropherograms and DNA alignment. Each sequence was processed the same way, corrections were made conservatively and tend to equalize non-variable sites but leave ambiguous calls in informative ones. Many sequences required extensive editing; in most cases the sequencing artefacts were systematic issues, caused by PCR errors and low template concentrations. In most cases sequences were obtained using reverse and forward primers, instead of enforcing the creation of a consensus contig assembly, we manually merged both senses during the correction step. The process of dataset assembly was iteratively repeated as new sequences were added.

### Extent of the multilocus dataset

We assembled a wide phylogenetic dataset including 824 lichen specimens and five sanger-sequenced nuclear loci (Tables S2-S4). Most specimens correspond to saxicolous species collected on limestone substrates across Southern Europe (Table S1, Figure S1). A subset of genetically divergent Central Asian and North American species to provide a broader phylogenetic context. The use of custom primers (Table S2) allowed us to assemble a 98% complete phylogenetic dataset. To deal with ambiguous basecalling caused by having partly dikaryotic specimens, the dataset was phased and concatenated into a data matrix including 910 phased haplotypes.

### Taxonomic credibility

After the datasets are assembled and aligned, their taxonomic consistency is assessed by several means. First, sequences of each locus are blasted against a local copy of the NCBI nt database (downloaded 3.2017), the resulting text output is processed in MEGAN v5 (*153*, *154*) to obtain an estimate of the Least common ancestor for each sequence. Samples containing foreign sequences or with a dubious adscription were excluded from the dataset. Because of the different representation of the loci in the NCBI database, the ITS, mtLSU and BT sequences are identified at least at a subfamily level (Caloplacoideae), the mcm7 sequences could be assigned to belong to the OSLEUM clade, while Efα, RPB1 and RPB2 could not be filtered taxonomically, and we relied on phylogenetic reconstructions for cleaning foreign sequences.

### Phasing of mixed sequences

The base calling of sanger sequences when using difficult material is often unsatisfactory, and often results in sequences containing numerous ambiguous sites. It is common practice in phylogenetic surveys, to retain ambiguous sites in the dataset when the overall sequence quality is high, since they have little influence on the topology, especially when used in a concatenated data matrix.

Base calling, being an automated process, does not take into account the process causing an ambiguous signal (e.g. primer degradation, presence of homopolymers, heterozygotic samples, paralogs, etc.). In datasets containing both intra and interspecific variability, like the one used, a single isolate may contain multiple variants in haploid or dikaryotic hyphae. This means that ambiguous sites accumulate in phylogenetically informative sites, significantly degrading the interpretation of the data. To avoid this type of dataset erosion we decided to curate sequences individually and carry out context-based haplotype inferences.

Upon the reception of sequence data, 1) forward and reverse sequences of the same specimen sequences were added and aligned to the data matrix in Geneious v7 using a fast MAFFT algorithm. 2) Using the electropherograms as context, low quality regions were eliminated and only high-quality fragments were used. 3) When high quality fragments did not cover the whole length of the sequence, they were repeated and appended to the alignment. 4) When forward and reverse sequences were properly overlapping and the context did not suggest a chimeric sequence was being created, all sequences were manually collapsed into one consensus. 5) Some samples however showed either different forward and reverse sequences or more often ambiguities reflecting a mixture of sequences similar to those existing in the dataset. In this case more than one sequence per sample was left in the alignment and the curated sequences were used to call phased haplotypes.

Most loci contained few mixed sequences and haplotypes could be phased manually. Four samples of ITS were mixed as well as two of RPB1, 12 of EF-α and 21 samples of β-tubulin. The mating type loci had also some mixed samples, 12 in matα and five in hmg. The nuclear mcm7 locus, however showed 80 samples which were clearly mixed and difficult to phase manually.

The genome assemblies showed no duplication of the mcm7 locus in *P. erodens*, in the assembly or looking at the patterns of Heterozygosity obtained by read-mapping. Haplotype phasing was carried out using the algorithm implemented in the program Phase v 2.1 (*155*–*158*); the input/output interface between alignment files and Phase was provided by fastphase (*159*).

To incorporate phased sequences into the population analyses, individual specimens with duplicated sequences in one locus were duplicated in all other datasets for population analyses. Haplotype phased specimens were treated as separate haploid entities and not as diploid in downstream population inferences. A total of 86 samples are duplicated in the final data matrix.

### Pylogenetic reconstructions, consensus and hybridization networks

Phylogenetic reconstruction on the duplicated dataset were carried out using iqtree (*160*) for Maximum Likelihood reconstructions and BEAST v2.2 (*161*) for Bayesian reconstructions. Initial selection of substitution models and partitions was carried out using the model testing capabilities of IQtree (*162*). The ML topologies of the five somatic loci were summarized using a majority rule consensus calculated in RaxML (*163*, *164*). Dendroscope v3 (*165*) was used to explore alternative consensus algorithms. Consensus networks captured best the phylogenetic incongruence between loci, we show the cluster network consensus and k-levels network. Pairwise hybridization networks were also used an hybridization number was estimated.

Bayesian reconstructions were carried out as a previous step for bGMYC (*166*), so a constant size coalescent model was used for each locus, which represents a conservative approach for the further the GMYC analysis. A strict clock and the substitution models selected in IQtree were also used.

### Species delimitation

To estimate the number of putative phylogenetic species based on single loci we used a multitree implementation of the General Mixed Yule Coalescent method as implemented in R package bGMYC(*166*). We chose a subset of 50 equally spaced trees from the posterior tree distribution of a Bayesian analysis using a constant size coalescent tree model. For each tree 500 alternative delimitations were obtained using a mcmc of 50K samples after burn-in of 40K iterations. A larger number of trees could not be used due to the large size of the dataset. The median number of species across all 2500 replicates is provided (Table S3). A consensus assignment was obtained using after summarizing the alternative bGMYC solutions as a coassignment matrix. This was converted to a dissimilarity matrix and processed using an iterative k-medoid(*167*) clustering method as implemented in function *pamk* of R package *fpc* (*168*). We imposed a search between 1 and 100 clusters and used the default average silhouette width criterion for the selection of the optimal number of groups. For the sake of visualization single locus clusters were also calculated using a maximum value of K of 10 or 15 clusters (Figures S2–S12). Additionally, we used single and multiple threshold GMYC(*55*) analyses for comparison, carried out using the consensus topology of the Bayesian analyses. Finally simpler OTU delimitations using the distance based ABGD(*56*) and ASAP(*169*) methods are provided (Table S2).

Using multilocus methods of species delimitation and discovery such as Tr2(*170*), SpedeStem (*171*) and Stacey (*172*) was the original aim of the survey. These require that species are either not connected through geneflow or need a putative delimitation to be used as starting point, which in our case is not readily attainable.

A multilocus interpretation of the bGMYC analyses is provided instead, making use of the obtained coassignment matrices. Each matrix summarizes the delimitation across multiple trees and replicates per locus as the probability that each pair of tips are assigned to the same cluster/putative species. Assuming that loci are not linked, and the species assignments are independent, the probability that two tips are coassigned across loci can be summarized two ways, as the average coassignment probability (average) or as the probability that two tips are coassigned in one or another locus, using the polynomial extension of p(A or B)=p(a) + p(b) -p(ab). The resulting pooled coassignment matrix was converted to a dissimilarity matrix (1-x) and clustered using a k-medoids method as implemented in function pamk of package fpc. Selection of k was automated using an average silhouette width as criterion.

### Inference of gene pools

Gene pools were estimated using BAPS(*125*). All loci were imported as concatenated alignments, duplicated sequences were included as independent haploid entities and not as diploid/polyploid data. Multilocus mixture clustering was carried iteratively using 20, 30, 50, 60, 80, 100, 150, 200 and 220 as maximum number of clusters. All analyses using a codon linkage model converged in identifying 10 genepools, using a less adequate linear linkage model, ten and eleven were equally likely solutions. The contribution of the different clusters under an Admixture model was calculated in an *a posteriori* run based on the precomputed mixture clusters. Finally the dataset was deduplicated in R. The proportions belonging to admixture fractions were recalculated, and the assignment to mixture clusters reinterpreted from the largest Admixture proportion. This deduplication did not alter the assignment to mixture clusters, but changed the admixture proportions.

The association between morphospecies and gene-clusters was surveyed using contingency tables and chi-square and statistical coindependence tests as implemented in R package vcd (*173*, *174*). The geographic distribution of cluster assignment was mapped using R packages ggplot2 (*175*), rnaturalearth and rnaturalearthdata(*176*).

### Estimating geneflow between gene pools

Geneflow between genepools was estimated using migrate-n v. 3.6.11 (*177*–*180*) Sampled specimens were clustered according to the assignment to mixture gene-pools estimated in BAPS, the connection matrix was imposed to be complete and initial values of theta and M were estimated from Fst table. A Bayesian estimation method was imposed using four incrementally heated chains (1,1.5,3,10K) and no swapping as it was intended for model selection. Each long chain is composed of 10M iterations in 10 replicate chains each with a burnin of 1M generations and 1M generations sampled every 100^th^ step. The results were tabulated and plotted in R using package igraph (CITE). The analyses were run usimg mutationally scaled population size and migration rate parameters. To interpret the data M was multiplied by the immigrated population size to obtain the number of migrants per generation 2Nem. We considered a conservative 2 migrants per generation as a significance threshold for plotting.

### Targeting MAT loci

In addition to the phylogenetically informative loci mentioned above, we developed specific primers for the mating type gene idiomorphs (MAT) using in the genome draft assemblies. To identify the MAT region we used a similar BLAST approach, but we included as target sequences both MAT 1.1.1 and MAT 1.1.2 genes as well as the flanking proteins SLA2 and APN2 from other Lecanoromycetes species. We identified all assemblies to be heterothallic and found no evidence of having both idiomorphs present in different parts of the genome (homoeothallic or pseudohomoeothallic). Heterothallism was further confirmed by mapping the raw reads back to the MAT regions of each idiomorph. Specific primers were developed to cover informative fractions of the MAT.1.1.1 or MATα gene as well as the gene of the HMG MAT 1.1.2 gene. Newly developed primers are summarized in (ST_2 primers).

### Phylogenetic diversity and cophylogenetic analysis of MAT genes

Nucleotide sequences of the two MAT idiomorphs (alpha and hmg) were used in the same way as the phylogenetic loci. Cophylogenetic analyses of mating type genes was carried out using two closely related methodologies aimed at interpreting cophylogenetic patterns in large data matrices implemented in R packages Paco and Random Tapas (*128*, *129*). Both methods largely rely on using the Gini coefficient (*181*, *182*)to measure the inequality among the values of the residual frequency distribution obtained using using procrustean superimposition of the host and symbiont matrices in Euclidean space. A Gini coefficient of 0 expresses perfect equality, where all values are the same, while a Gini coefficient of 1 (or 100%) expresses maximal inequality among values. To run cophylogenetic analyses on such a highly diverse dataset, required collapsing the nucleotide dataset. To simplify the analyses and retain the distinction between nucleotide and aminoacid datasets, idiomorph alignments were collapsed to OTUs using a conservative ABGD run, resulted in a matrix containing 243 alpha and 225 hmg OTUs respectively. Phylogenetic ML trees were pruned to include a representative sequence per distance cluster.

### Network analysis of MAT idiomorph distribution

Nucleotide sequences of the two MAT idiomorphs (alpha and hmg) were translated to aminoacid sequences in Geneiousv7, using the alignment to a complete genomic reference as reading frame reference. Aminoacid sequence alignments were imported into R using package phangorn, where they were collapsed into haplotypes. Haplotype datasets were collapsed to 99, 97 and 95% similarity clusters using Cd-HIT (*133*, *183*, *184*). The resulting cluster assignments were imported into R, where the occurrence of idiomorph haplogroups per specimen were tabulated.

The table of idiomorph association at individual level was used to generate a unipartite graph using igraph v.1.3.2 (*132*). To identify reproductive compartments in the network, we first partitioned it identifying its maximal connected components (i.e. compartments). Further, modularity was estimated using four alternative algorithms: walktrap (*185*), louvain(*134*), labelProp(*186*) and infomap(*187*). Alternative partitions were validated and compared using the methodology implemented in R package *ROBin* (*188*). Alternative partitions are compared in terms of their stability against increasing levels of random perturbation, measured using the Variation of Information metric (VI) (*189*). Model comparison was carried out using the Gaussian Process ranking method for time series implemented in package *geprege* (*190*). A key limitation of the used methodology is its dependence on the adequacy of the randomized networks used as null models. We explored two alternative randomization approaches based on the observed network. The configuration model (CM) implemented in random function of package Robin generates a random graph with the same degree sequence as the original but with a randomized group structure. Meanwhile the program RandNetGen (*136*) was used to generate a random graph based on the dk-series 2.1 model, which preserves the joint degree distribution and the clustering coefficient of the original network but not the full clustering spectrum.

Alternatively, the idiomorph table was processed as a bipartite network using package bipartite (*131*, *191*). Network compartmentalization was estimated using function *compart*, and modularity was surveyed using function *computeModules*. The network structure and the differences between levels were calculated using functions *networklevel* and *classlevel*.

### Genomic sequencing assembly and annotation

The genome of *Pyrenodesmia erodens* was sequenced using material from axenic culture; all other genomes have been assembled from metagenomic libraries of complete thallus fragments. Library construction and sequencing were outsourced to the High-Throughput Genomics and Bioinformatic Analysis center of the Huntsman Cancer Institute of the University of Utah (Salt Lake City, UT). The reference genome of *P. erodens* was assembled using two libraries, a paired-end sequencing library constructed using the TruSeq DNA PCR-Free sample preparation kit (Illumina) with an insert size of 550bp, and a mate-pair library constructed using Nextera Mate Pair Library Preparation Kit (Illumina) sequenced using three target insert sizes 3Kb, 5.3Kb and 10Kb.

For the metagenomic libraries we used either TruSeq DNA PCR-free (Illumina) or nano kits (Illumina) depending of the DNA yield of the siolation process in any case using a more conservative 350bp insert size. All libraries were sequenced using a HiSeq 125 cycle paired-end sequencing v4 protocol using two lanes of an Illumina Hiseq-2000 platform.

The quality of the resulting libraries was quantified using fastqc (*192*). They were quality trimmed and filtered using trimmomatic (*193*), and assembled using spades v 3.6.2 (*194*). Quality and completeness were assessed using quast (*195*) and busco (*196*). For *P. erodens* we evaluated missassemblies graphically by mapping mate-pair libraries onto the assembled genome. We identified several problematic regions, especially around large homopolymer repeats (polyC). These were manually split and rescaffolded using sspace v2(*197*).

Lichen metagenomes were assembled using spades v 3.6.2 (*194*) using the algorithm for isolates. Additional metagenome assemblies, metaspades and megahit were used for cross-validation. Metagenome assemblers were used but discarded as their intended use is beyond that of this manuscript. algorithms were also employed, but were ultimately discarded. Although several alternative metagenome assembly algorithms were tried, isolate but in addition alternative approaches were also metagenome assemblers. Fungal scaffolds were identified using a custom strategy implemented in R (SUPPLEMENT). The graphic strategy akin to that used in a blobology approach makes use of k-mer coverage, length and GC content to plot the sequencing distribution of scaffolds. Decision on the origin of each assembled scaffold was made using sequence comparison strategies and different sources of evidence. A nucleotide blast was carried out against the reference genome (*P.erodens*) using blastn (*198*); diamond (*199*) was used for nucleotide to protein alignments (blastx) using a dataset of publically available Lecanoromycete proteins (*Xanthoria parietina, Cladonia uncialis* and *P. erodens*) as well as against a copy of the ncbi’s nr dataset (DATE). The blast results were analyzed using megan v5 (*200*) to obtain a taxonomic assignment. Finally the presence of Lecanoromycete single copy orthologs was evaluated using busco (*201*). The different lines of evidence were concatenated in a binary string, including whether the nucleotide blast was positive or not, whether the blastx was positive or not and if any of the hits were the best hits for a particular reference protein in the database, whether the scaffold was identified as fungal or Lecanoromycete and whether Lecanoromycete buscos were identified. The process continued onto gene prediction and annotation as described below. After a first round of gene prediction and functional annotation, genomes were additionally filtered to eliminate minor scaffolds containing only proteins identified as being of bacterial origin using the eggNOG (*202*) database and eggNOG-mapper (*203*).

Gene prediction and annotation of the *P.erodens* genome differs from the rest of the genomes. In general we used the funannotate v(1.7.4) (*204*) as the main analytical pipeline to mask the genome, carry out gene prediction and annotation. Because we had no RNA evidence to train gene models, gene prediction was purely based in preexisting protein databases, RefSeq (*205*) and the specific lichen protein database used for filtering including the proteins identified in the *P. erodens* genome. The latter genomic draft was initially annotated using the Maker v3 (*206*) pipeline and and Bast2GO (*207*). A first set of proteins was used as an additional source of evidence for the final gene prediction, which used the funannotate pipeline (*204*) to keep coherent with the rest of the survey. Differently to the proposed strategy implemented in funannotate we opted to carry out Eggnogmapper (*203*) Interproscan (*208*) Antismash (*209*) certain analyses independently and later integrate them into funannotate. Funannotate compare was used to obtain a first tabulate of genome features across the dataset. Antismash results were further explored to find Biosynthetic Genes using similarity clustering as implemented in bigScape (*210*). Synteny was explored using multiple collinearity criteria as implemented in MCScanX (*211*). Ripping was surveyed using the TheRipper online tool (*212*). The presence of repetitive and mobile elements in the unmasked genomes was carried out using Repeatmasker/modeler (*213*) and the last publicly available repbase database (*214*). Additionally, protein motifs found associated with giant mobile elements (*215*) were searched individually using blast (*149*)alignments.

### Phylogenomic inference

Predicted protein sets were processed in Orthofinder (*216*) which was used to identify Orthogroups and single copy orthologs (OGs). The OG aminoacid alignments were used to query the corresponding mRNA sets, using gene names, which were put together using a simple R script. Nucleotide alignments pero OG were then carried out using mafft (*151*) in Auto mode and trimmed using trimal (*217*).

Single gene trees were calculated in iqtree (*218*) using bootstrap replicates to obtain nodal support values. ML topologies were compiled into a single file and processed in raxml (*163*) to generate a consensus and calculate internode centrainty and tree certainty support values (IC/TC) (*219*, *220*). A cluster network consensus was estimated in dendroscope (*165*), where pairwise hybridization networks (*121*) And hybridization numbers were also calculated.

### Data exploration in R

Genome annotation data were imported into R, explored and visualized using functions implemented in the karyoploteR (*221*) and GenomicRanges (*222*) packages for both visualization and summary. The genomic dataset was split into three range sets. First, telomeric and subtelomeric regions were taken out. We chose as subtelomeric the terminal 250Kb of scaffolds where telomeric repeats were identified. Second LRARs not contained within telomeric regions were taken out. They were extended to cover an additional 30Kb on each flank, overlapping ranges were merged. The remaining ranges were considered as the general coding fraction. The density of genomic features was estimated using function *kpPlotDensity* and a window size of 50Kb. Differences in density across sliding windows between genomic were visualized using violin plots in *ggplot2*. Differences between regions were tested using pairwise Wilcoxon rank sum test with continuity correction. The loss of genes within syntenic blocks was obtained by parsing the output of MCScanX. The overall influence of the density of mobile elements on the estimated gene loss was modelled using *glm* Poisson regression. Further visualizations were carried out using package *circlize* (*223*).

### Reproducibility and data availability

All datasets, scripts and supplementary materials are organized as Rmarkdown documents https://github.com/ferninfm/pyrenodesmia_phylogenomics and https://github.com/ferninfm/pyrenodesmia_phylogenetics. They are provided as html and pdf files but can also be compiled for reproducibility. Sanger-sequenced data has been made available through NCBIs genebank under accession numbers XXXXX-XXXXX. Short read files and assembled genomic drafts are made available under Bioproject #XXXXXX and Biosamples #XXXXXX #XXXXXX.

## Notes

https://github.com/ferninfm/pyrenodesmia_phylogenomics

### Competing Interest Statement

The authors have declared no competing interest.

