## Supplementary_material for "Introgressive Descent and Hypersexuality Drive The Evolution Of Sexual Parasitism and Morphological Reduction In a Fungal Species Complex"

Table S1. Sampling localities

| Loc | lon | lat | alt | Description | country |
|---|---|---|---|---|---|
| 1 | -104,8 | 31,89 | 1874,2 | Culberson County, Texas, 79847, United States | USA |
| 2 | -6,002869 | 43,058258 | 1601,43 | Pista de Ricabo al Alto de Ventana, Teverga, Asturias / Asturies, 24144, España | Spain |
| 3 | -5,923793 | 43,187917 | 1768,46 | Quirós, Asturias / Asturies, España | Spain |
| 4 | -5,92205 | 43,184632 | 1702,85 | Quirós, Asturias / Asturies, 33217, España | Spain |
| 5 | -5,914669 | 43,173728 | 1520,92 | Quirós, Asturias / Asturies, 33217, España | Spain |
| 6 | -5,900369 | 43,166979 | 1344,28 | Despegue de Peral, Carretera de Bárzana a Pola de Lena, Lena, Asturias / Asturies, España | Spain |
| 7 | -5,880573 | 42,915947 | 1088,57 | LE-3418, Sena de Luna, León, Castilla y León, España | Spain |
| 8 | -5,3935 | 36,752488 | 0 | Sendero Puerto de las Presillas, Grazalema, Sierra de Cádiz, Cádiz, Andalucía, 11610, España | Spain |
| 9 | -5,361113 | 36,758342 | 746,82 | A-372, Grazalema, Sierra de Cádiz, Cádiz, Andalucía, 11610, España | Spain |
| 10 | -5,236414 | 43,42929 | 881,65 | Fuente Mergullines, Camino de Cofiño a El Bustaco, Cofiñu, Parres, Asturias / Asturies, 33343, España | Spain |
| 11 | -5,197068 | 36,493066 | NA | Estepona, Costa del Sol Occidental, Málaga, Andalucía, España | Spain |
| 12 | -5,180057 | 36,693849 | NA | Rezume de la Puentezuela, Carril de Ronda, Ronda, Serranía de Ronda, Málaga, Andalucía, España | Spain |
| 13 | -5,029904 | 36,795446 | NA | Cueva de los Covarones, A-366, Ronda, Serranía de Ronda, Málaga, Andalucía, España | Spain |
| 14 | -5,010227 | 36,797477 | NA | A-366, Ronda, Serranía de Ronda, Málaga, Andalucía, España | Spain |
| 15 | -4,985969 | 36,781801 | NA | Mirador del Guarda Forestal, A-366, El Burgo, Sierra de las Nieves, Málaga, Andalucía, España | Spain |
| 16 | -4,90789 | 36,805358 | NA | MA-5401, El Burgo, Sierra de las Nieves, Málaga, Andalucía, España | Spain |
| 17 | -4,834034 | 36,802974 | NA | MA-5401, Casarabonela, Sierra de las Nieves, Málaga, Andalucía, 29551, España | Spain |
| 18 | -4,531894 | 36,958022 | 1113,08 | MA-9016, Antequera, Málaga, Andalucía, España | Spain |
| 19 | -4,174057 | 37,113849 | 1439,93 | Carril de Alcantar, Loja, Comarca de Loja, Granada, Andalucía, 18313, España | Spain |
| 20 | -3,649337 | 41,547543 | 940,29 | SG-V-9322, Valdevacas de Montejo, Segovia, Castilla y León, 40542, España | Spain |
| 21 | -3,648698 | 36,825398 | 1085,21 | Los Guájares, Comarca de la Costa Granadina, Granada, Andalucía, 18615, España | Spain |
| 22 | -3,642661 | 41,540422 | 998,69 | SG-V-9322, Valdevacas de Montejo, Segovia, Castilla y León, 40542, España | Spain |
| 23 | -3,624054 | 36,84452 | 599,5 | Guájar Alto, Los Guájares, Comarca de la Costa Granadina, Granada, Andalucía, 18615, España | Spain |
| 24 | -3,519666 | 36,824156 | 239,49 | Carretera de Vélez de Benaudalla a Motril, Vélez de Benaudalla, Comarca de la Costa Granadina, Granada, Andalucía, 18670, España | Spain |

| | | | | | |
|---|---|---|---|---|---|
| **25** | -3,461392 | 41,908045 | 1126,06 | BU-910 (antigua BU-911), Espinosa de Cervera, Burgos, Castilla y León, España | Spain |
| **26** | -3,447873 | 40,885559 | 886,45 | Camino de Servicio del Canal del Jarama CYII, Patones, Sierra Norte, Comunidad de Madrid, España | Spain |
| **27** | -3,438029 | 37,119329 | 2048,44 | Carretera de Granada a Sierra Nevada, Güéjar Sierra, Comarca de la Vega de Granada, Granada, Andalucía, 18160, España | Spain |
| **28** | -3,432251 | 37,123926 | 2116,21 | Monte Ahí de Cara: Vista Norte, Carretera de Granada a Sierra Nevada, Güéjar Sierra, Comarca de la Vega de Granada, Granada, Andalucía, 18196, España | Spain |
| **29** | -3,423803 | 37,115763 | 2182,3 | Camino Real de los Neveros, Güéjar Sierra, Comarca de la Vega de Granada, Granada, Andalucía, 18196, España | Spain |
| **30** | -3,391818 | 36,83482 | 1681,46 | El Hoyo de las Encinas, Órgiva, Comarca de la Alpujarra Granadina, Granada, Andalucía, España | Spain |
| **31** | -3,391227 | 36,835709 | 1658,39 | El Hoyo de las Encinas, Órgiva, Comarca de la Alpujarra Granadina, Granada, Andalucía, España | Spain |
| **32** | -3,389633 | 36,836094 | 1625,7 | El Hoyo de las Encinas, Órgiva, Comarca de la Alpujarra Granadina, Granada, Andalucía, España | Spain |
| **33** | -3,389188 | 36,834965 | 1617,29 | El Hoyo de las Encinas, Órgiva, Comarca de la Alpujarra Granadina, Granada, Andalucía, España | Spain |
| **34** | -3,388603 | 36,832327 | 1591,82 | El Hoyo de las Encinas, Órgiva, Comarca de la Alpujarra Granadina, Granada, Andalucía, España | Spain |
| **35** | -3,386622 | 41,955963 | 1066,94 | BU-910, Carazo, Burgos, Castilla y León, 09610, España | Spain |
| **36** | -3,383445 | 36,831329 | 1344,76 | El Alhayón, Órgiva, Comarca de la Alpujarra Granadina, Granada, Andalucía, España | Spain |
| **37** | -3,374267 | 41,963748 | 1091,69 | BU-910, Carazo, Burgos, Castilla y León, 09610, España | Spain |
| **38** | -2,864949 | 40,943863 | 832,86 | CM-1003, Bujalaro, Guadalajara, Castilla-La Mancha, 19245, España | Spain |
| **39** | -2,615909 | 41,015161 | 1059,25 | GU-118, Sigüenza, Guadalajara, Castilla-La Mancha, 19266, España | Spain |
| **40** | -2,615288 | 41,010722 | 1088,57 | GU-118, Sigüenza, Guadalajara, Castilla-La Mancha, 19268, España | Spain |
| **41** | -2,515461 | 37,244481 | 1904,96 | de Serón a Gérgal, El Cortijuelo, Bacares, Valle del Almanzora, Almería, Andalucía, España | Spain |
| **42** | -2,007678 | 36,853595 | NA | Aparcamiento Playa El Playazo, Camino al Playazo, La Ermita, Níjar, Almería, Andalucía, 04115, España | Spain |
| **43** | -1,460845 | 37,505958 | 337,55 | RM-D20, El Cantal, Lorca, Alto Guadalentín, Región de Murcia, España | Spain |
| **44** | -1,15035 | 37,551927 | 369,27 | El Castillo, Diputación de Perín, Cartagena, Campo de Cartagena y Mar Menor, Región de Murcia, España | Spain |
| **45** | -1,113029 | 37,612046 | 285,64 | RM-E26, Perín, Diputación de Perín, Cartagena, Campo de Cartagena y Mar Menor, Región de Murcia, España | Spain |
| **46** | -1,012201 | 40,117537 | 1866,03 | A-2520, Camarena de la Sierra, Gúdar-Javalambre, Teruel, Aragón, España | Spain |
| **47** | -1,001859 | 40,129439 | 1761,97 | A-2520, La Puebla de Valverde, Gúdar-Javalambre, Teruel, Aragón, España | Spain |
| **48** | -0,934862 | 40,427901 | 1581,72 | A-226, Corbalán, Comunidad de Teruel, Teruel, Aragón, España | Spain |
| **49** | -0,710011 | 40,342699 | 1552,64 | Carretera Alcalá, Mora de Rubielos, Gúdar-Javalambre, Teruel, Aragón, 44431, España | Spain |
| **50** | -0,648093 | 40,388817 | 1923,47 | A-2705, Valdelinares, Gúdar-Javalambre, Teruel, Aragón, España | Spain |
| **51** | -0,647311 | 40,387089 | 1906,89 | A-2705, Valdelinares, Gúdar-Javalambre, Teruel, Aragón, España | Spain |
| **52** | -0,636361 | 40,387081 | 1904,24 | El Villarejo, Valdelinares, Gúdar-Javalambre, Teruel, Aragón, España | Spain |
| **53** | -0,324966 | 38,650215 | 1006,62 | Alcoleja, el Comtat, Alacant / Alicante, Comunitat Valenciana, 03814, España | Spain |
| **54** | -0,0616 | 38,563007 | 193,59 | Camí de la Cantera, L'Albir, l'Alfàs del Pi, la Marina Baixa, Alacant / Alicante, Comunitat Valenciana, 03581, España | Spain |
| **55** | 4,82664 | 43,747536 | 151,53 | Piste des Lombards, Les Baux-de-Provence, Arles, Bouches-du-Rhône, Provence-Alpes-Côte d'Azur, France métropolitaine, 13520, France | France |
| **56** | 5,2773 | 44,172724 | 1892 | Restaurant Le Vendran, Route du Mont-Ventoux, Bédoin, Carpentras, Vaucluse, Provence-Alpes-Côte d'Azur, France métropolitaine, 84410, France | France |
| **57** | 5,959118 | 44,697475 | 1498,57 | Le Barry, Route du Col du Noyer, Saint-Étienne-en-Dévoluy, Le Dévoluy, Gap, Hautes-Alpes, Provence-Alpes-Côte d'Azur, France métropolitaine, 05250, France | France |

| | | | | | |
|---|---|---|---|---|---|
| **58** | 5,981688 | 44,698628 | 1608,88 | Route du Col du Noyer, Saint-Étienne-en-Dévoluy, Le Dévoluy, Gap, Hautes-Alpes, Provence-Alpes-Côte d'Azur, France métropolitaine, 05250, France | France |
| **59** | 5,985443 | 44,692332 | 1667,76 | D 17, Le Noyer, Gap, Hautes-Alpes, Provence-Alpes-Côte d'Azur, France métropolitaine, 05500, France | France |
| **60** | 6,197296 | 46,479921 | 1135,43 | Le Bugnonet, Bassins, District de Nyon, Vaud, 1269, Schweiz/Suisse/Svizzera/Svizra | Switzerland |
| **61** | 6,465392 | 45,991996 | 1755 | Acces Falaise du Col de la Colombière, Le Grand-Bornand, Annecy, Haute-Savoie, Auvergne-Rhône-Alpes, France métropolitaine, 74450, France | France |
| **62** | 7,638056 | 44,06722 | NA | D 43, La Brigue, Nice, Alpes-Maritimes, Provence-Alpes-Côte d'Azur, France métropolitaine, 06430, France | France |
| **63** | 10,984768 | 45,682396 | 1352,21 | Strada Provinciale 14 dell'Alta Valpantena, Villa, Erbezzo, Verona, Veneto, 37020, Italia | Italy |
| **64** | 11,137556 | 45,881957 | 890,78 | Strada vecia di Serrada, Dieneri, Piazza, Terragnolo, Comunità della Vallagarina, Provincia di Trento, Trentino-Alto Adige/Südtirol, 38064, Italia | Italy |
| **65** | 13,706009 | 46,594166 | 1755,96 | Dobratsch-Gipfelweg, Bad Bleiberg, Bezirk Villach-Land, Kärnten, 9530, Österreich | Austria |
| **66** | 13,878713 | 45,622263 | 421,18 | Riserva Naturale Val Rosandra / Naravni rezervat Dolina Glinščice, Località Basovizza, Altipiano Est, Basovizza / Bazovica, Trieste, Friuli-Venezia Giulia, 34018, Italia | Italy |
| **67** | 13,881852 | 45,551369 | 394,02 | Utrdba San Sergio nad Črnim Kalom, Črni Kal, Katinara, Črni Kal, Koper / Capodistria, Upravna enota Koper / Unità amministrativa Capodistria, 6275, Slovenija | Slovenia |
| **68** | 14,03333 | 42,06667 | NA | Strada Statale 487 di Caramanico Terme, Pacentro, L'Aquila, Abruzzo, Italia | Italy |
| **69** | 14,080214 | 45,787402 | 751,87 | Nad votlo steno, Strane, Postojna, 6225, Slovenija | Slovenia |
| **70** | 14,13028 | 49,92722 | NA | národní přírodní rezervace Koda, Ekísova úniková, Korno, okres Beroun, Střední Čechy, 267 27, Česko | Czech Republic |
| **71** | 14,20194 | 45,28556 | NA | Poletište Vojak, R,P,O,, Grad Opatija, Primorsko-goranska županija, Hrvatska | Croatia |
| **72** | 14,21083 | 45,30222 | NA | R,P,O,, Poklon, Vela Učka, Grad Opatija, Primorsko-goranska županija, Hrvatska | Croatia |
| **73** | 14,30722 | 49,99472 | NA | ev,18, Ořech, okres Praha-západ, Střední Čechy, 252 25, Česko | Czech Republic |
| **74** | 14,33556 | 49,98917 | NA | národní přírodní památka Černé rokle, V Borovičkách, Kosoř, okres Praha-západ, Střední Čechy, 153 00, Česko | Czech Republic |
| **75** | 14,46333 | 45,58194 | NA | Ilirska Bistrica, Loška Dolina, Slovenija | Slovenia |
| **76** | 15,40111 | 47,33111 | NA | Zustiegsweg, Fladnitz an der Teichalm, Bezirk Weiz, Steiermark, 8163, Österreich | Austria |
| **77** | 15,42361 | 47,3625 | NA | Eiserne-Kling Hütte, Wöllingergrabenweg, Lantsch, Breitenau am Hochlantsch, Bezirk Bruck-Mürzzuschlag, Steiermark, 8614, Österreich | Austria |
| **78** | 15,42361 | 47,3625 | NA | Eiserne-Kling Hütte, Wöllingergrabenweg, Lantsch, Breitenau am Hochlantsch, Bezirk Bruck-Mürzzuschlag, Steiermark, 8614, Österreich | Austria |
| **79** | 15,42361 | 47,3625 | NA | Eiserne-Kling Hütte, Wöllingergrabenweg, Lantsch, Breitenau am Hochlantsch, Bezirk Bruck-Mürzzuschlag, Steiermark, 8614, Österreich | Austria |
| **80** | 19,05028 | 43,09667 | NA | Žabljak - Trsa - Plužine, Pošćenski katun, Pašina voda, Opština Žabljak, 84220, Crna Gora / Црна Гора | Montenegro |
| **81** | 19,05111 | 43,09833 | NA | Sedlo pass, Žabljak - Trsa - Plužine, Pošćenski katun, Pašina voda, Opština Žabljak, 84220, Crna Gora / Црна Гора | Montenegro |
| **82** | 19,05222 | 43,09806 | NA | Žabljak - Trsa - Plužine, Pošćenski katun, Pašina voda, Opština Žabljak, 84220, Crna Gora / Црна Гора | Montenegro |
| **83** | 19,78167 | 42,51306 | NA | Dolina Grebaje, Škala, Opština Gusinje, Crna Gora / Црна Гора | Montenegro |
| **84** | 20,0125 | 42,68111 | NA | Lijepi Do, Katun Lijepi do, Opština Plav, Crna Gora / Црна Гора | Montenegro |
| **85** | 20,01306 | 42,68 | NA | Lijepi Do, Katun Lijepi do, Opština Plav, Crna Gora / Црна Гора | Montenegro |
| **86** | 20,015 | 42,68222 | NA | Lijepi Do, Katun Lijepi do, Opština Plav, Crna Gora / Црна Гора | Montenegro |
| **87** | 20,015 | 42,68222 | NA | Lijepi Do, Katun Lijepi do, Opština Plav, Crna Gora / Црна Гора | Montenegro |
| **88** | 20,015 | 42,68222 | NA | Lijepi Do, Katun Lijepi do, Opština Plav, Crna Gora / Црна Гора | Montenegro |
| **89** | 20,08833 | 42,58861 | NA | Stanet e Bjeshkëve te Belegut, Komuna e Deçanit / Opština Dečane, 51000, Kosova / Kosovo | Kosovo |

| | | | | | |
|---|---|---|---|---|---|
| **90** | 20,13917 | 42,75722 | NA | Hajla, Opština Rožaje, Crna Gora / Црна Гора | Montenegro |
| **91** | 20,13917 | 42,75722 | NA | Hajla, Opština Rožaje, Crna Gora / Црна Гора | Montenegro |
| **92** | 20,64556 | 41,94889 | 1560 | Restelicë, Komuna e Dragashit / Opština Dragaš, Kosova / Kosovo | Kosovo |
| **93** | 20,65 | 40,03333 | NA | Αετόπετρα, Δήμος Κόνιτσας, Περιφερειακή Ενότητα Ιωαννίνων, Περιφέρεια Ηπείρου, Αποκεντρωμένη Διοίκηση Ηπείρου - Δυτικής Μακεδονίας, 441 00, Ελλάς | Greece |
| **94** | 20,71871 | 39,96811 | NA | Άγιος Βλάσσιος, Αρίστης - Πάπιγκου, Πάπιγκο, Δήμος Ζαγορίου, Περιφερειακή Ενότητα Ιωαννίνων, Περιφέρεια Ηπείρου, Αποκεντρωμένη Διοίκηση Ηπείρου - Δυτικής Μακεδονίας, 440 16, Ελλάς | Greece |
| **95** | 20,71871 | 39,96811 | NA | Άγιος Βλάσσιος, Αρίστης - Πάπιγκου, Πάπιγκο, Δήμος Ζαγορίου, Περιφερειακή Ενότητα Ιωαννίνων, Περιφέρεια Ηπείρου, Αποκεντρωμένη Διοίκηση Ηπείρου - Δυτικής Μακεδονίας, 440 16, Ελλάς | Greece |
| **96** | 20,71871 | 39,96811 | NA | Άγιος Βλάσσιος, Αρίστης - Πάπιγκου, Πάπιγκο, Δήμος Ζαγορίου, Περιφερειακή Ενότητα Ιωαννίνων, Περιφέρεια Ηπείρου, Αποκεντρωμένη Διοίκηση Ηπείρου - Δυτικής Μακεδονίας, 440 16, Ελλάς | Greece |
| **97** | 20,76833 | 39,96972 | NA | Μικρό Πάπιγκο, Δήμος Ζαγορίου, Περιφερειακή Ενότητα Ιωαννίνων, Περιφέρεια Ηπείρου, Αποκεντρωμένη Διοίκηση Ηπείρου - Δυτικής Μακεδονίας, 440 04, Ελλάς | Greece |
| **98** | 20,77028 | 39,98083 | NA | Ο3, Δήμος Ζαγορίου, Περιφερειακή Ενότητα Ιωαννίνων, Περιφέρεια Ηπείρου, Αποκεντρωμένη Διοίκηση Ηπείρου - Δυτικής Μακεδονίας, Ελλάς | Greece |
| **99** | 20,779883 | 40,346289 | NA | Δήμος Κόνιτσας, Περιφερειακή Ενότητα Ιωαννίνων, Περιφέρεια Ηπείρου, Αποκεντρωμένη Διοίκηση Ηπείρου - Δυτικής Μακεδονίας, Ελλάς | Greece |
| **100** | 20,78639 | 39,99444 | NA | Δρακόλιμνη, Νταβάλιστα, Δήμος Ζαγορίου, Περιφερειακή Ενότητα Ιωαννίνων, Περιφέρεια Ηπείρου, Αποκεντρωμένη Διοίκηση Ηπείρου - Δυτικής Μακεδονίας, Ελλάς | Greece |
| **101** | 20,78639 | 39,99444 | NA | Δρακόλιμνη, Νταβάλιστα, Δήμος Ζαγορίου, Περιφερειακή Ενότητα Ιωαννίνων, Περιφέρεια Ηπείρου, Αποκεντρωμένη Διοίκηση Ηπείρου - Δυτικής Μακεδονίας, Ελλάς | Greece |
| **102** | 20,80861 | 39,98806 | NA | Σάδι Μύγας, Δήμος Ζαγορίου, Περιφερειακή Ενότητα Ιωαννίνων, Περιφέρεια Ηπείρου, Αποκεντρωμένη Διοίκηση Ηπείρου - Δυτικής Μακεδονίας, Ελλάς | Greece |
| **103** | 25,78333 | 35 | NA | 'Ιεράπετρας - Σητείας, Κοινότητα Ιεράπετρας, Δημοτική Ενότητα Ιεράπετρας, Δήμος Ιεράπετρας, Περιφερειακή Ενότητα Λασιθίου, Περιφέρεια Κρήτης, Αποκεντρωμένη Διοίκηση Κρήτης, 722 00, Ελλάς | Greece |
| **104** | 25,78333 | 35 | NA | 'Ιεράπετρας - Σητείας, Κοινότητα Ιεράπετρας, Δημοτική Ενότητα Ιεράπετρας, Δήμος Ιεράπετρας, Περιφερειακή Ενότητα Λασιθίου, Περιφέρεια Κρήτης, Αποκεντρωμένη Διοίκηση Κρήτης, 722 00, Ελλάς | Greece |
| **105** | 25,9 | 39,15 | NA | Σκάλα Ερεσού, Δήμος Δυτικής Λέσβου, Περιφερειακή Ενότητα Λέσβου, Περιφέρεια Βόρειου Αιγαίου, Αποκεντρωμένη Διοίκηση Αιγαίου, Ελλάς | Greece |
| **106** | 27,33417 | 40,75583 | NA | Mürefte Barbaros Yolu, Uçmakdere Mahallesi, Şarköy, Tekirdağ, Marmara Bölgesi, Türkiye | Greece |
| **107** | 33,88444 | 44,40444 | NA | Ифигения, Заросшая тупиковая лестница, Ялтинский городской совет, Автономна Республіка Крим, 298687, Україна | Ukraine |
| **108** | 35,58111 | 38,70639 | NA | Salih Gıdık Caddesi, Tavlusun Mahallesi, Kayseri, Melikgazi, Kayseri, İç Anadolu Bölgesi, 38165, Türkiye | Turkey |
| **109** | 45,39778 | 37,18528 | NA | بالستان, بخش مرکزی, شهرستان ارومیه, استان آذربایجان غربی, ایران | Iran |
| **110** | 45,45028 | 37,92528 | NA | دهستان جزیره, بخش ایلخچی, شهرستان اسکو, استان آذربایجان شرقی, ایران | Iran |
| **111** | 51,72889 | 43,58389 | NA | Мунайлинский район - Мұнайлы ауданы, Маңғыстау облысы, 130600, Қазақстан | Kazakhstan |
| **112** | 52,05528 | 44,24306 | NA | Тиген, Мангистауский район - Маңғыстау ауданы, Маңғыстау облысы, 130400, Қазақстан | Kazakhstan |
| **113** | 52,06611 | 44,21333 | NA | Тиген, Мангистауский район - Маңғыстау ауданы, Маңғыстау облысы, 130400, Қазақстан | Kazakhstan |
| **114** | 52,09778 | 43,95 | NA | Мангистауский район - Маңғыстау ауданы, Маңғыстау облысы, Қазақстан | Kazakhstan |
| **115** | 52,65278 | 44,15556 | NA | Бейнеу-Актау, Мангистауский район - Маңғыстау ауданы, Маңғыстау облысы, 130402, Қазақстан | Kazakhstan |

| | | | | | |
|---|---|---|---|---|---|
| **116** | 54,99889 | 45,02389 | NA | река Манаши, Бейнеуский район - Бейнеу ауданы, Маңғыстау облысы, Қазақстан | Kazakhstan |
| **117** | 55,28 | 51,94111 | NA | Дачная улица, Известковое, Краснокоммунарский поссовет, Сакмарский район, Оренбургская область, Приволжский федеральный округ, Россия | Russia |
| **118** | 56,09806 | 53,55278 | NA | Торатау, крутая каменистая тропа, Урман-Бишкадакский сельсовет, Ишимбайский район, Башкортостан, Приволжский федеральный округ, Россия | Russia |
| **119** | 57,66944 | 51,11167 | NA | Набережная улица, Айтуар, Кувандыкский городской округ, Оренбургская область, Приволжский федеральный округ, Россия | Russia |
| **120** | 57,77528 | 53,36083 | NA | Кулганинский сельсовет, Бурзянский район, Башкортостан, Приволжский федеральный округ, Россия | Russia |
| **121** | 58,77056 | 52,04556 | NA | Уральский сельсовет, Кваркенский район, Оренбургская область, Приволжский федеральный округ, Россия | Russia |
| **122** | 58,80028 | 52,08667 | NA | Уральский сельсовет, Кваркенский район, Оренбургская область, Приволжский федеральный округ, Россия | Russia |

**Figure S1.** Map of sampling localities

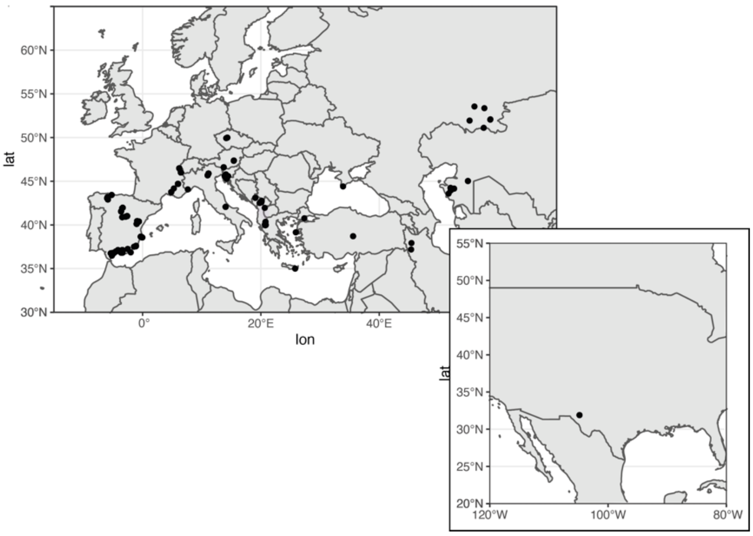

**Table S2.** Newly developed primer sequences, including additional ones for RPB2 and mtLSU which were sequenced but not used in the final dataset.

| *Primer Name* | *Target Gene* | *Motif* | *Tm* | *Ta* | *length* | *sequence length* |
|---|---|---|---|---|---|---|
| ITS_267f | Fungal ITS | GGTCAAACTTGGTCATTTAG | 56 | 51 | 20 | 992bp |
| ITS_1259r | Fungal ITS | CCGAAACATCCTCTACAAAT | 56 | 51 | 20 | - |
| tub_5100f | β-Tubulin | GGCAAATAGCTACAATGGTACCTC | 70 | 65 | 24 | 1015bp |
| tub_6115R | β-Tubulin | GGCCATCATGTTCTTTGGATCATA | 68 | 63 | 24 | - |
| EFA_713F | EFα | GTCACCGCGATTTCATCAAGA | 62 | 57 | 21 | 740bp |
| EFA_1453R | EFα | CCACGACGGATTTCCTTGAC | 62 | 57 | 20 | - |
| mcm7_4959F | MCM7 | CATCTCATCACAGTCCGTGGCATTG | 76 | 71 | 25 | 729bp |
| mcm7_5688R | MCM7 | CTCGTGGTGCAACCTTGGTRATGTA | 76 | 71 | 25 | - |
| RPB1_191F | RPB1 | ACCGTGGTATTAGGTGTGGGACTTG | 76 | 71 | 25 | 891bp |
| RPB1_1082R | RPB1 | TCCATGTAGGTCGCAACGTGGAATT | 74 | 69 | 25 | - |
| RPB2_A_1498F | RPB2 | TCCCAAGTGCTCAATCGCTA | 60 | 55 | 20 | 960bp |
| RPB2_A_2458R | RPB2 | CATGCGCTCATCGTAGTTCG | 62 | 57 | 20 | - |
| RPB2_B_2365F | RPB2 | CCCAGATCATAACCAGTCTCCTC | 70 | 65 | 23 | 954bp |
| RPB2_B_3319R | RPB2 | GTGGCCATTGTACATGACCTC | 63 | 58 | 21 | - |
| ml3a_alt | mtLSU | GCTGGTTTTCCGCGAAACCTATATAAG | 78 | 73 | 27 | 1050bp |
| mtlsu_7315R | mtLSU | GGCTCACCTTATCCCGAAGTTACGT | 75 | 70 | 25 | - |

**Table S3.** Description of the sequenced datasets. Parsimony informative sites and average diversities were calculated in mega v11. All values are calculated excluding outgroup sequences.

| | Phylogenetically informative genes | | | | | Mating type genes | | | |
|---|---|---|---|---|---|---|---|---|---|
| | ITS | β-tub | EFα | Mcm7 | RPB1 | α | hmg | α∪hmg | α∩hmg |
| Number of samples | 824 | 804 | 820 | 815 | 776 | 571 | 561 | 727 | 423 |
| Completeness | 100% | 98% | 99% | 99% | 94% | . | . | 88% | 51% |
| Number of sequences | 910 | 882 | 901 | 899 | 833 | 586 | 566 | . | . |
| Duplicated samples | 86 | 78 | 82 | 85 | 58 | 16 | 6 | . | . |
| Number of haplotypes | 515 | 428 | 448 | 486 | 337 | 389 | 321 | . | . |
| Translated haplotypes | . | . | . | . | . | 252 | 199 | . | . |
| 99% similarity peptids | . | . | . | . | . | 169 | 105 | . | . |
| of which connected | . | . | . | . | . | 136 | 91 | . | . |
| ABGD | 190 | 124 | 69 | 114 | 96 | 91 | 2(166) | . | . |
| ASAP best score | 368 | 338 | 218 | 378 | 157 | 161 | 2(224) | . | . |
| ASAP min | 288 | 239 | 217 | 204 | 56 | 117 | 2(78) | . | . |
| GMYCsingle clust/ent | 38/197 | 37/111 | 38/199 | 18/82 | 42/188 | 6/13 | 18/78 | . | . |
| GMYCmulti clust/ent | 6/23 | 54/223 | 39/195 | 30/109 | 45/121 | 56/382 | 46/205 | | |
| bGMYC pamk | 56 | 86 | 15 | 97 | 60 | . | . | | |
| bGMYc median | 34 | 32 | 5 | 50 | 24 | 16 | 37 | | |
| Alignment length | 712 | 626 | 426 | 633 | 660 | 946 | 848 | . | . |
| Variable sites | 392 | 251 | 179 | 325 | 310 | 509 | 468 | | |
| Parsimony Inf. sites | 327 | 236 | 164 | 282 | 269 | 379 | 369 | . | . |
| Nucleotide diversity | 0.07 | 0.07 | 0.05 | 0.06 | 0.05 | 0.06 | 0.05 | | |
| Haplotype diversity | 0.566 | 0.483 | 0.497 | 0.541 | 0.404 | 0.681 | 0.572 | | |

Table S4. Hardwired cluster distances between phylogenetic trees.

| | ITS | β-tub | EFα | Mcm7 | RPB1 |
|---|---|---|---|---|---|
| ITS | 0.00 | 848.00 | 838.00 | 849.00 | 834.00 |
| β-tub | 848.00 | 0.00 | 865.00 | 865.00 | 831.00 |
| EFα | 838.00 | 865.00 | 0.00 | 861.00 | 841.00 |
| Mcm7 | 849.00 | 865.00 | 861.00 | 0.00 | 858.00 |
| RPB1 | 834.00 | 831.00 | 841.00 | 858.00 | 0.00 |

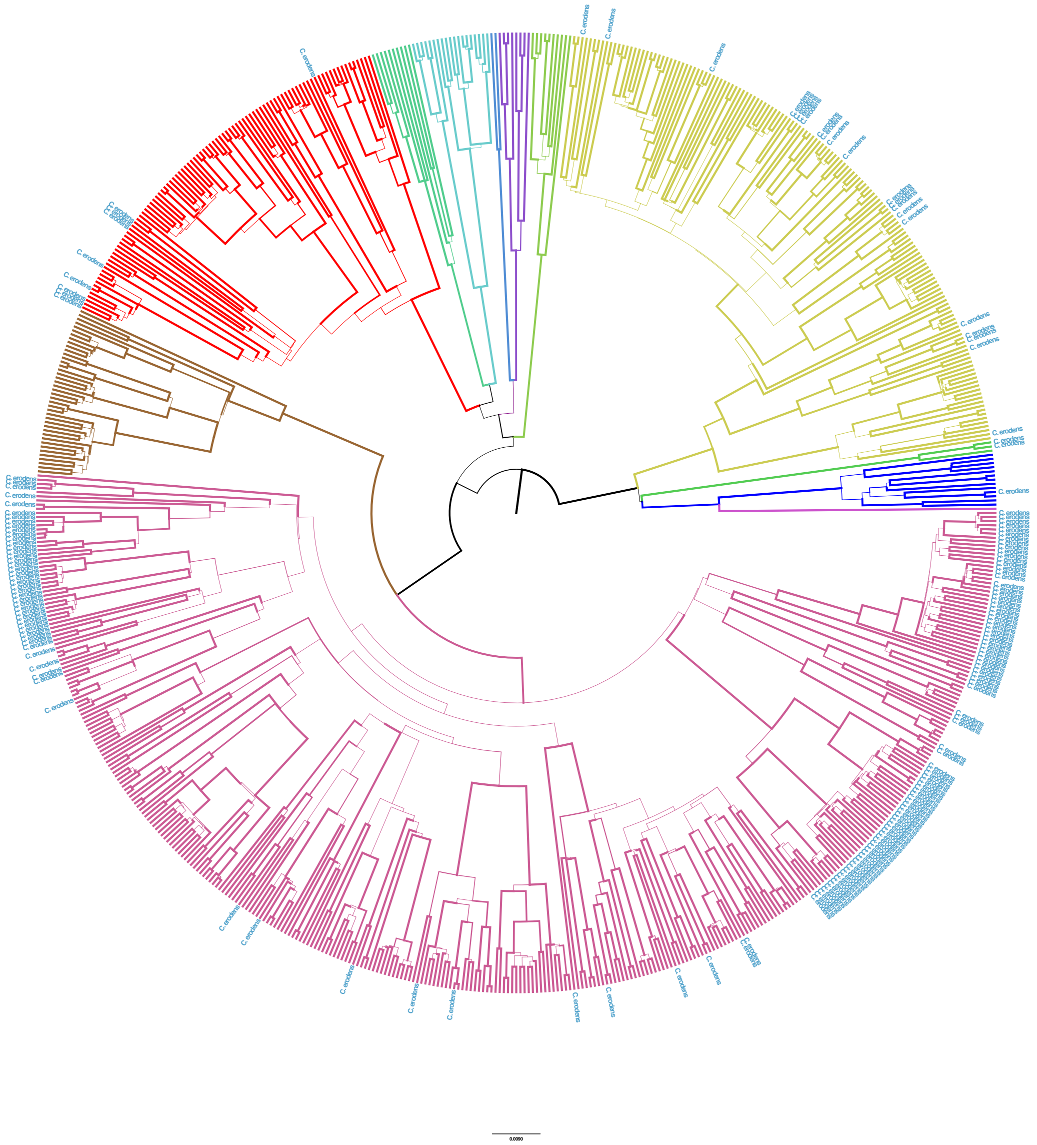

Figure S2. Single locus trees ITS

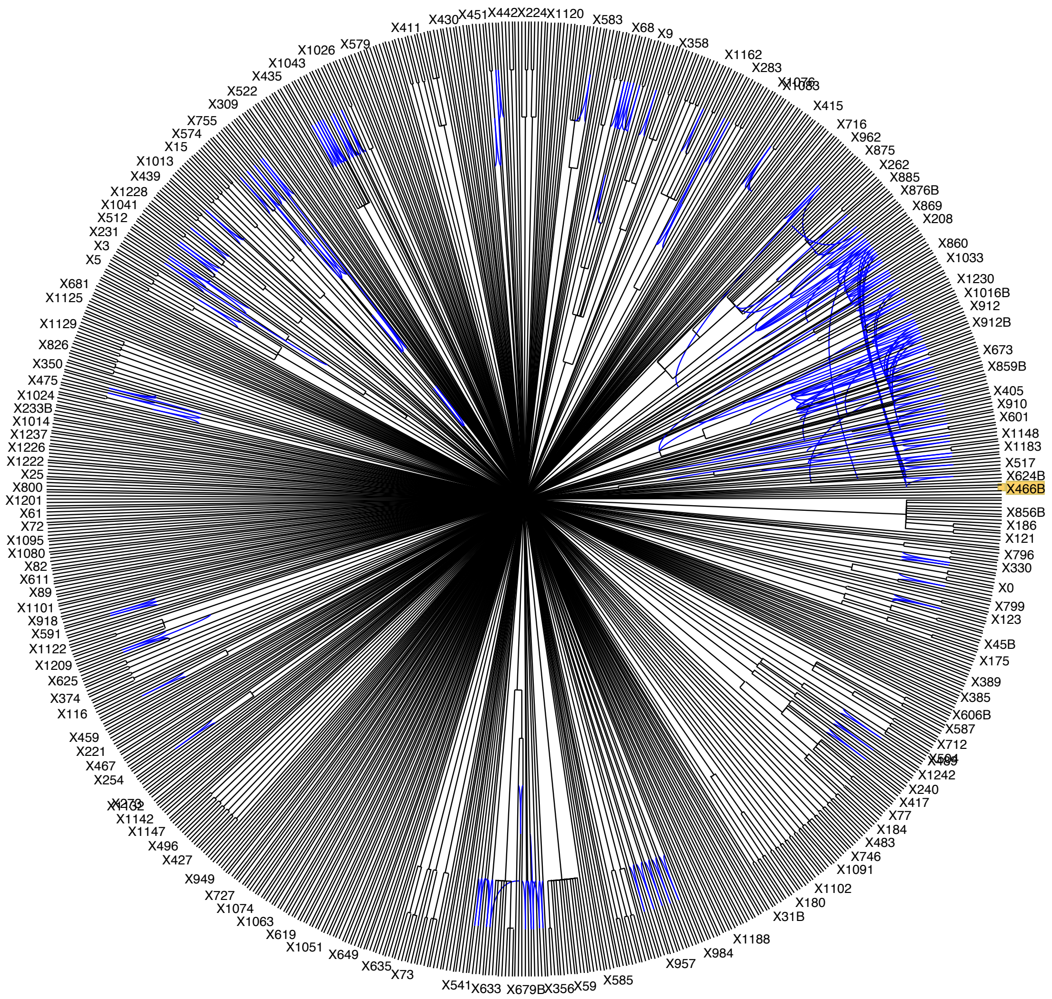


Figure S3. Consensus cluster network based on the ML phylogenies of ITS, mcm7, β-tubulin, EFα and RPB1.

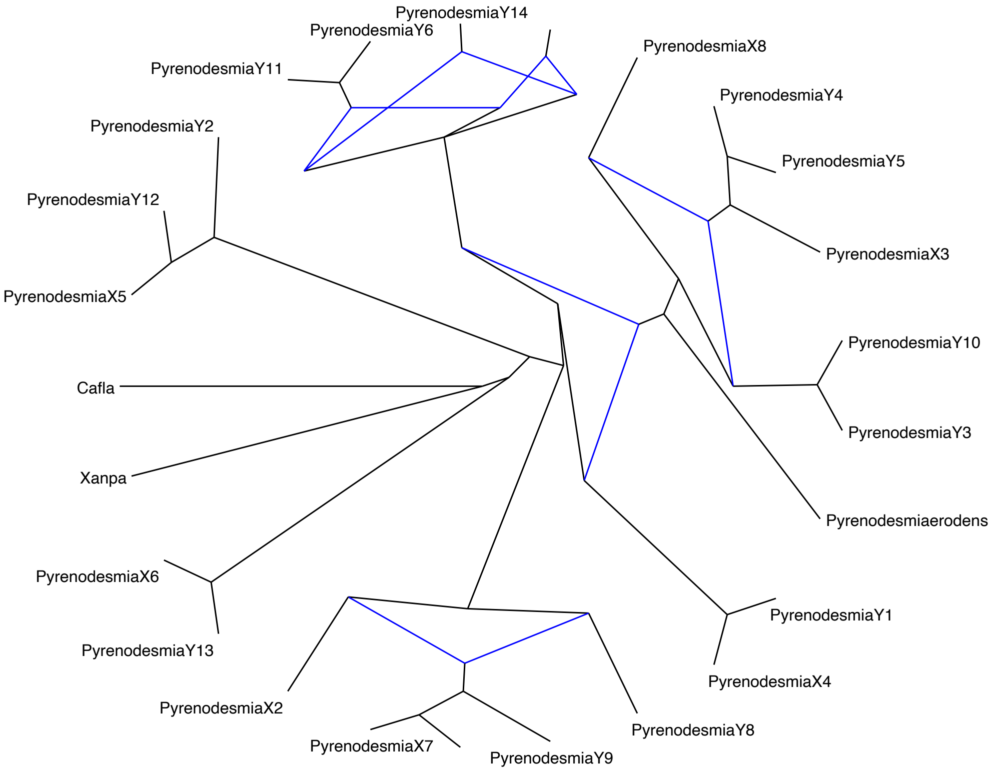


Figure S4. Phylogenomic consensus network based on 2767 ML reconstructions of single copy orthologs.

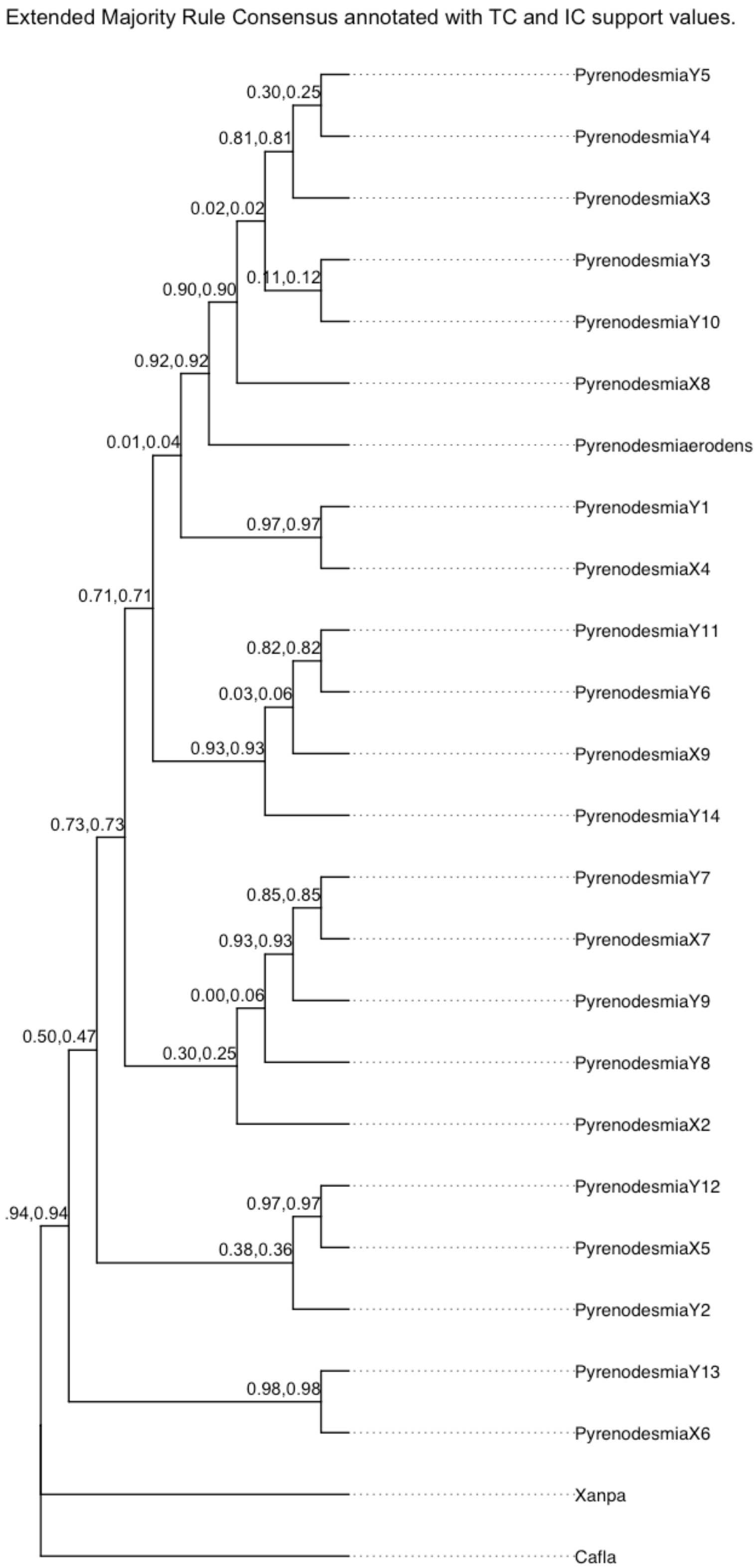


Figure S5. Phylogenomic consensus tree. Nodal support values represent Internode and Tree certainties.

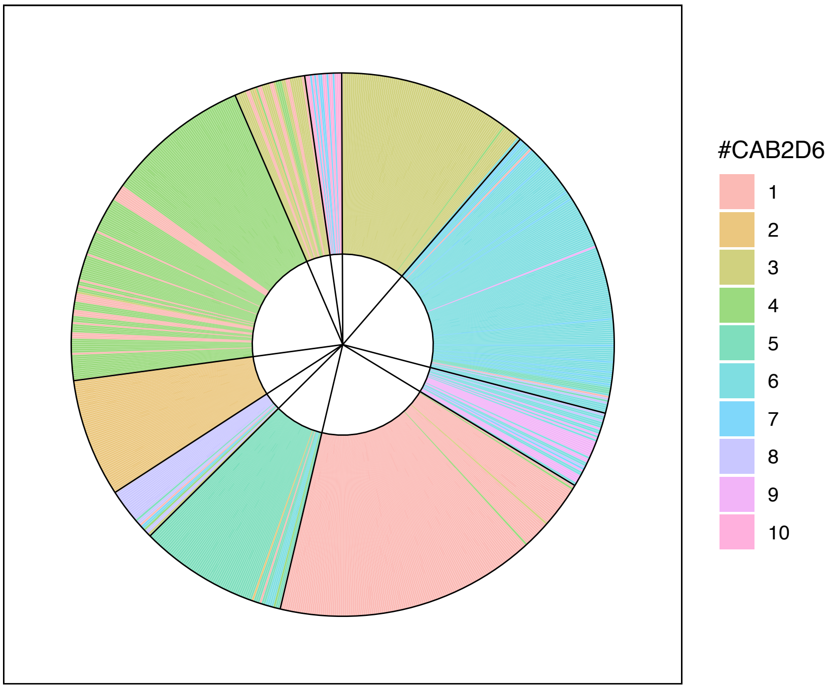


**FIGURE S6. Barplot of ITS bGMYC genetic clusters obtained imposing a small maximum number of clusters to pamk (k=10).** The sectors on the circle plot reflect the Assignment to BAPS multilocus clusters as shown across all figures.

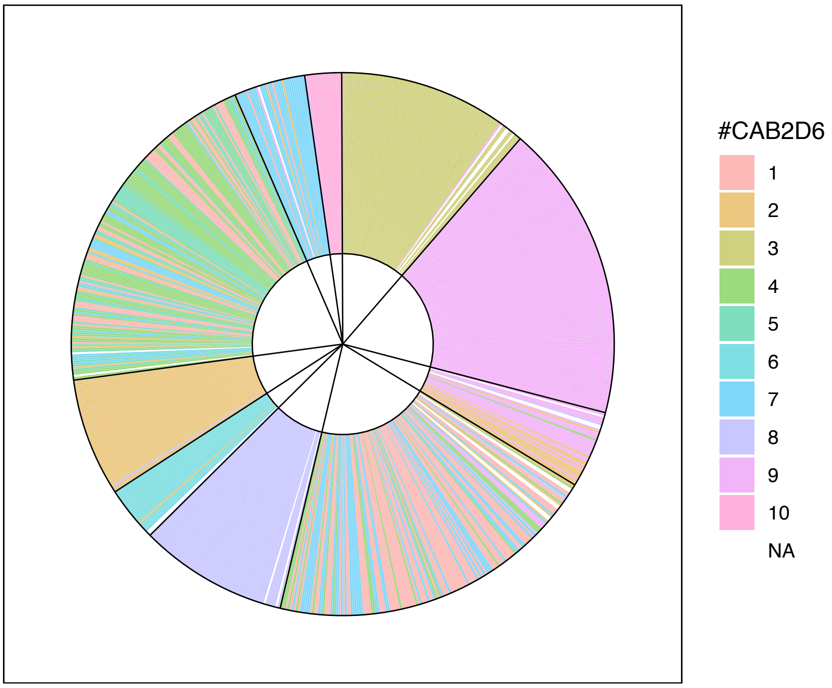


**FIGURE S7. Barplot of B-tubulin bGMYC genetic clusters** obtained imposing a small maximum number of clusters (k=10)

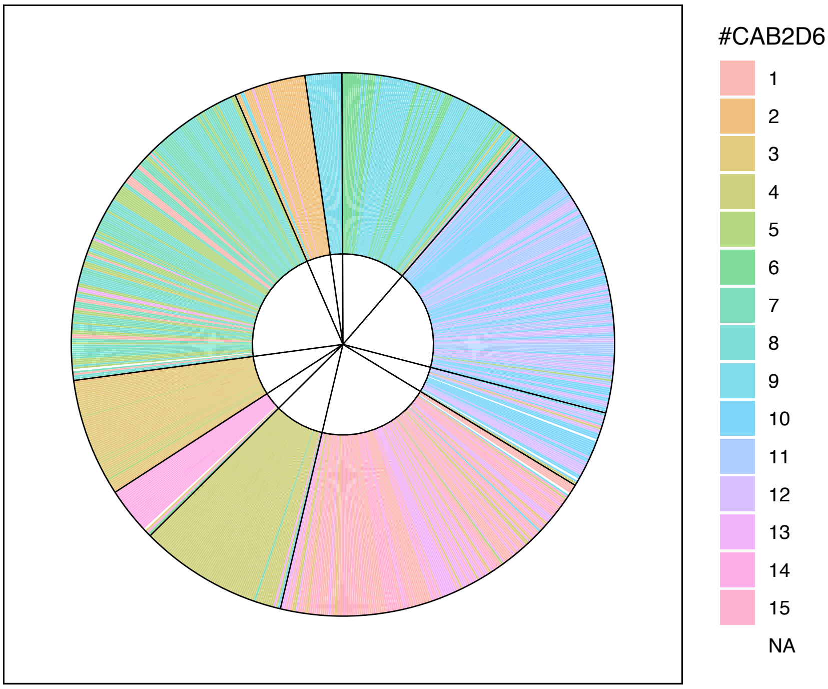


**FIGURE S8. Elongation Factor α bGMYC genetic clusters** obtained implementing the optimum number of clusters identified by pamk (k=15)

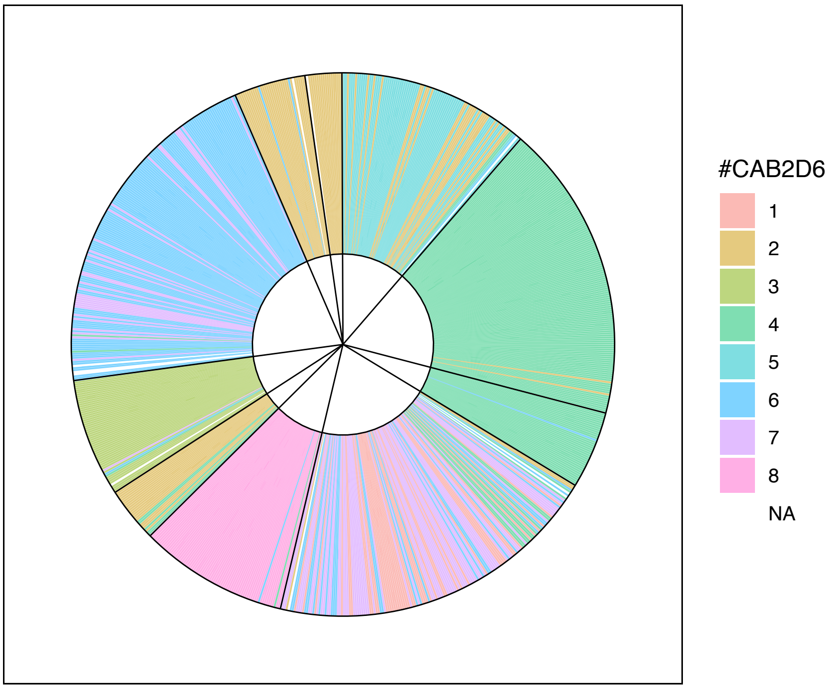


**FIGURE S9. Barplot of MCM7 bGMYC genetic clusters** obtained imposing a small maximum number of clusters (k=10)

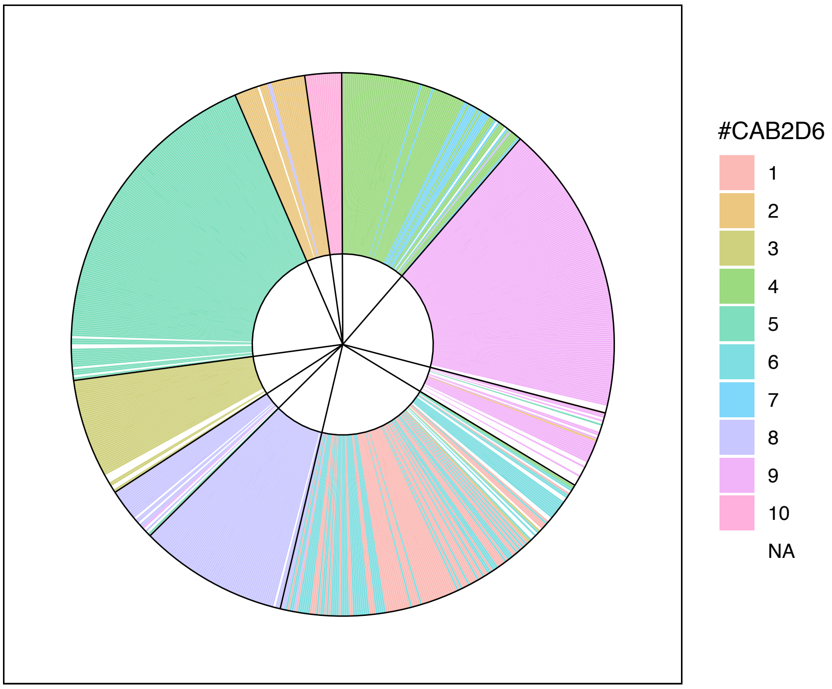


**FIGURE S10. Barplot of RPB1 bGMYC genetic clusters** obtained imposing a small maximum number of clusters (k=10/15)

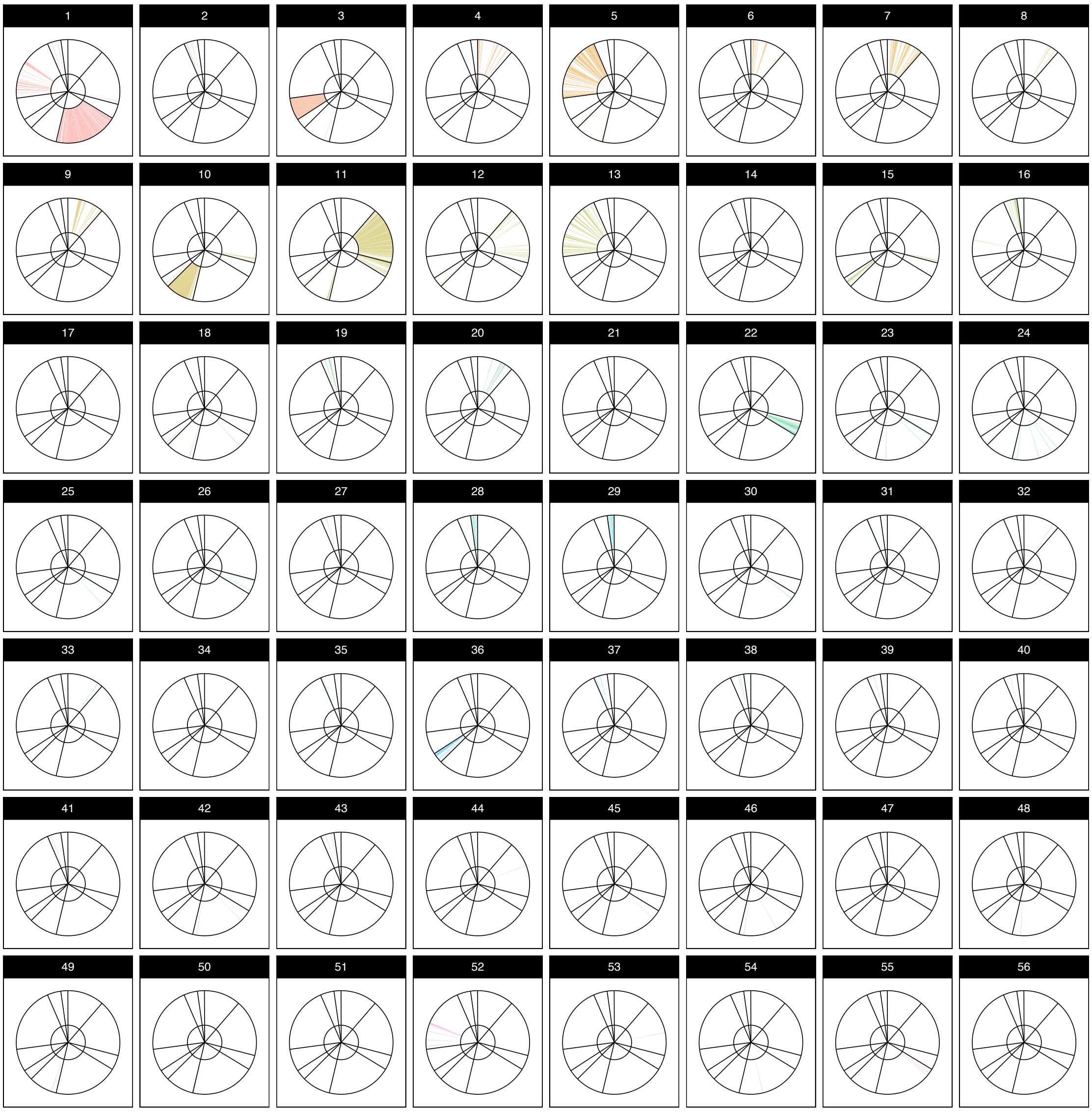


**FIGURE S11.** Barplots of ITS bGMYC genetic clusters using the optimum number of clusters as identified in pamk.

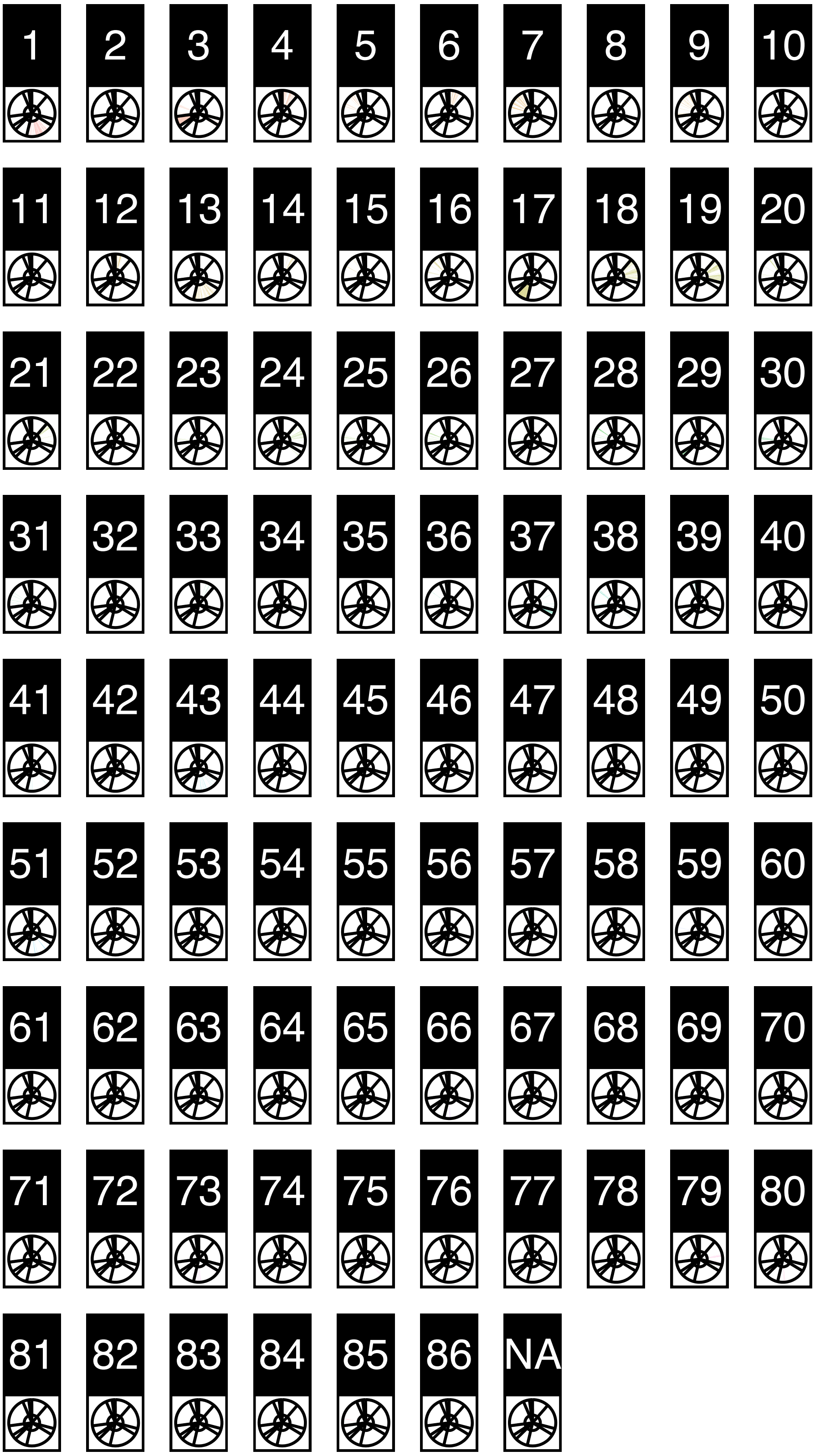


**FIGURE S12.** Barplots of B-tubulin bGMYC genetic clusters using the optimum number of clusters as identified in pamk.

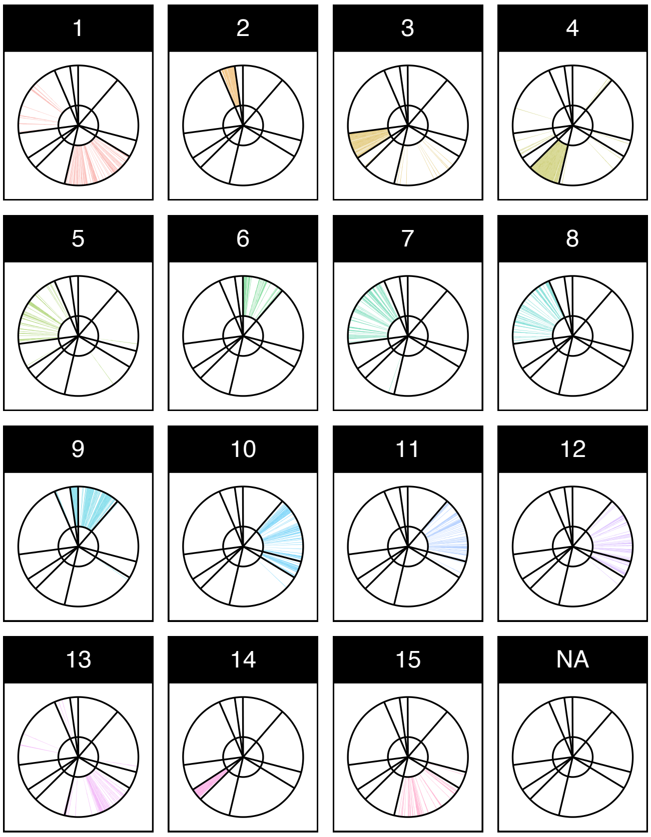


**FIGURE S13.** Barplot of EFa single locus bGMYC genetic clusters using the optimum number of clusters as identified in pamk.

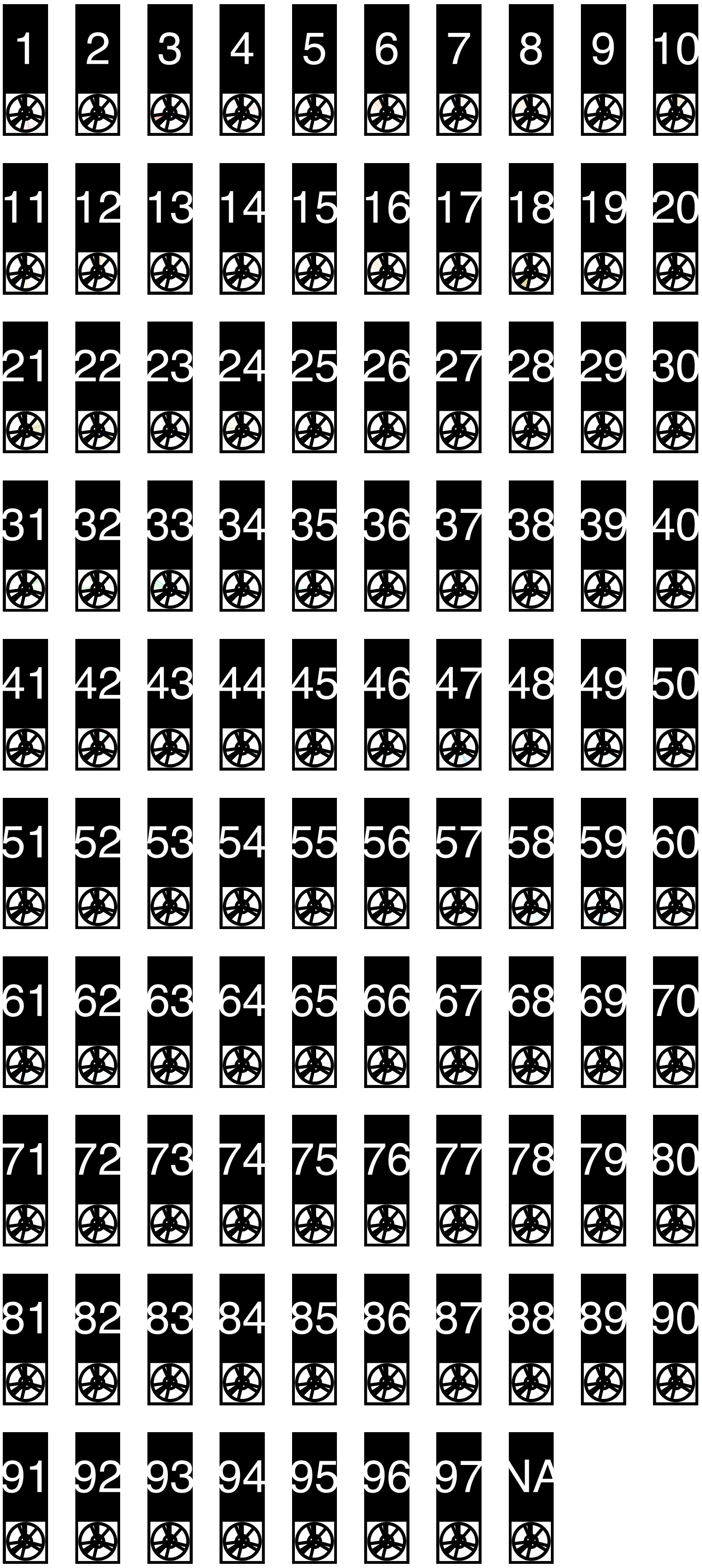


**FIGURE S14.** Barplot of MCM7 single locus bGMYC genetic clusters using the optimum number of clusters as identified in pamk.

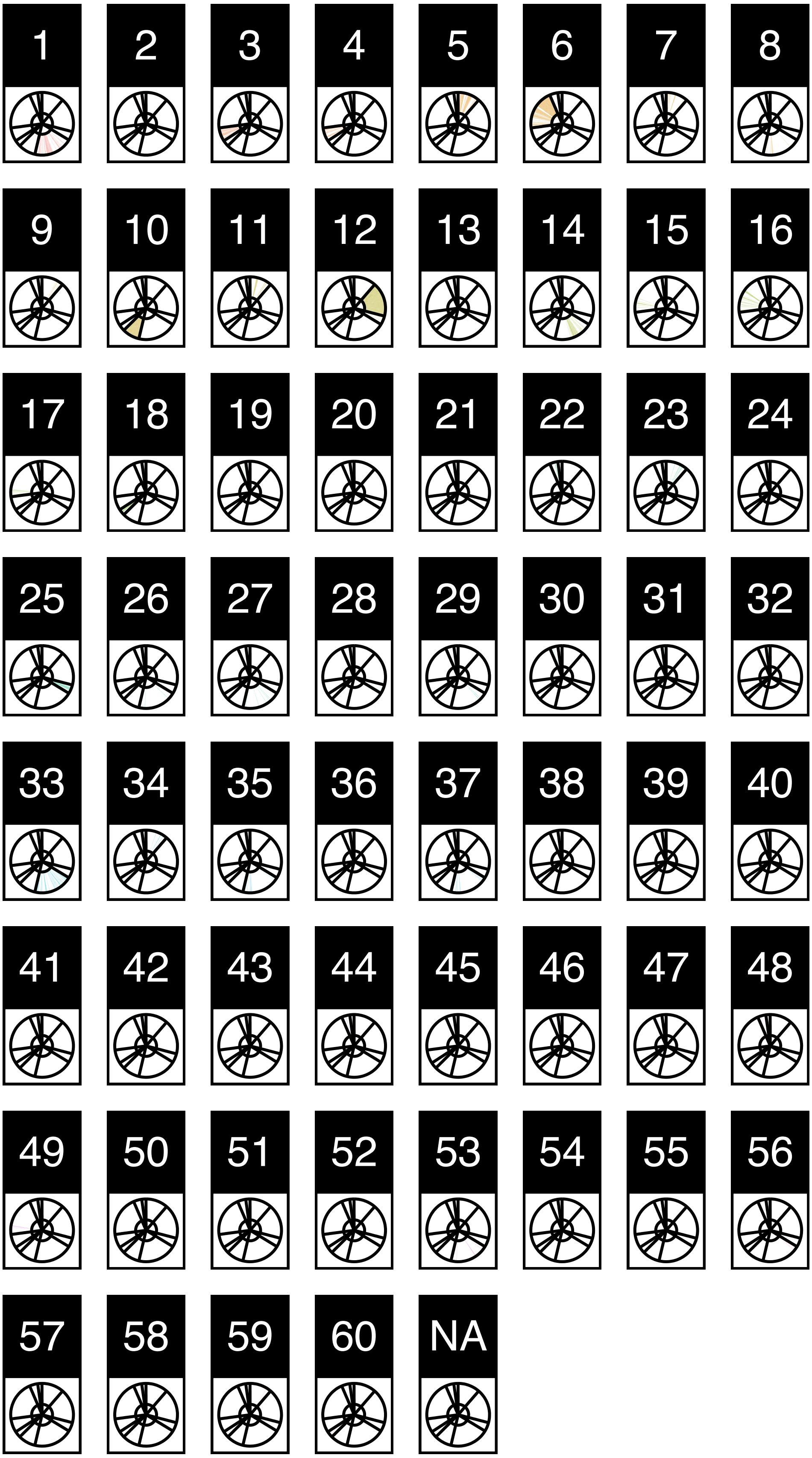


**FIGURE S15.** Barplot of RPB1 single locus bGMYC genetic clusters using the optimum number of clusters as identified in pamk.

**FIGURE S16.** Barplot of MCM7 single locus bGMYC genetic clusters using the optimum number of clusters as identified in pamk.

**FIGURE S17.** Single locus bGMYC genetic clusters obtained imposing a small maximum number of clusters (k=10/15)

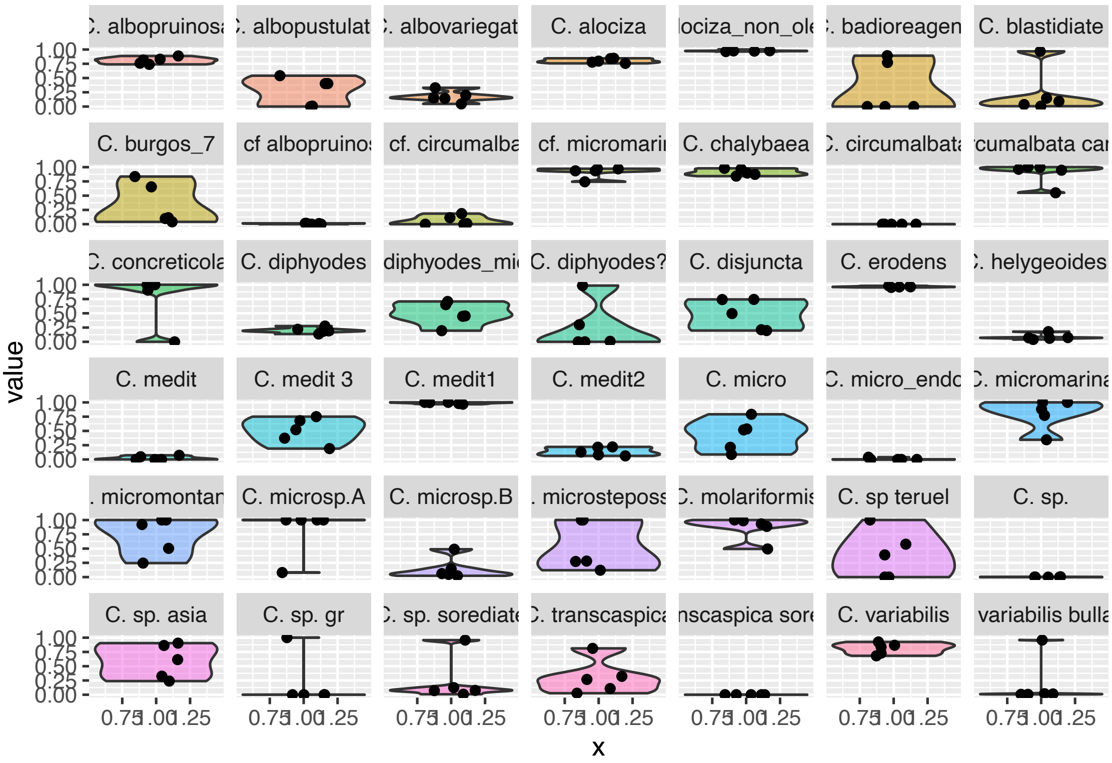


**FIGURE S18.** Phylogenetic concordance between phylogenetic reconstructions grouped at operational phenotypic units used across the survey.

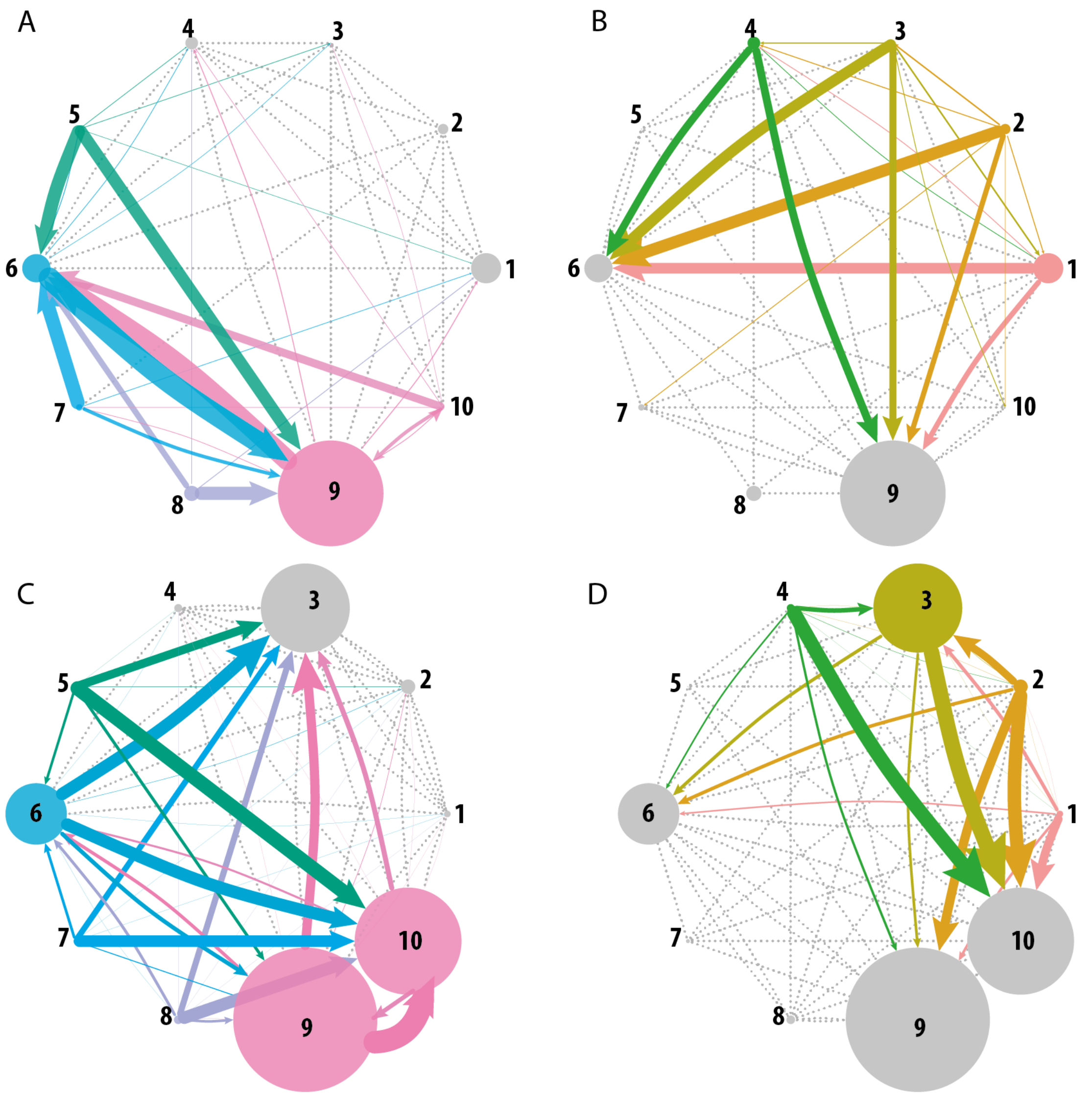


FIGURE S19. Migrate-n networks obtained for the individual mating type idiomorphs. a–b| Mating type alpha; c–d| Mating type hmg.

| Module | # nodes | # edges | density | centrality | Path length |
|---|---|---|---|---|---|
| Complete | 676 | 8203 | 0.036 | 0.059 | 3.654 |
| main | 601 | 8104 | **0.045** | **0.074** | **3.656** |
| 1 | 79 | 1124 | 0.365 | 0.092 | 1.825 |
| 2 | 19 | 84 | **0.491** | **0.402** | **1.754** |
| 4 | 39 | 271 | 0.366 | 0.287 | 2.0553 |
| 5 | 38 | 299 | **0.426** | **0.126** | **1.704** |
| 6 | 64 | 1105 | **0.548** | **0.084** | **1.583** |
| 7 | 45 | 188 | 0.190 | 0.463 | 3.351 |
| 9 | 23 | 132 | **0.522** | **0.121** | **1.668** |
| 11 | 58 | 247 | 0.149 | 0.135 | 2.715 |
| 12 | 55 | 979 | **0.659** | **0.146** | **1.395** |
| 14 | 51 | 355 | 0.278 | 0.373 | 2.307 |
| 15 | 74 | 2422 | 0.897 | 0.038 | 1.130 |
| 16 | 12 | 59 | 0.894 | 0.023 | 1.106 |
| 18 | 44 | 268 | 0.283 | 0.254 | 2.313 |
| secondary | 75 | 99 | - | - | - |
| 3 | 4 | 4 | 0.666 | 0.666 | 1.333 |
| 8 | 2 | 1 | 1 | - | 1 |
| 10 | 2 | 1 | 1 | - | 1 |
| 13 | 9 | 21 | 0.583 | 0.536 | 1.417 |
| 17 | 4 | 6 | 1 | 0 | 1 |
| 19 | 5 | 10 | 1 | 0 | 1 |
| 20 | 2 | 1 | 1 | - | 1 |
| 21 | 2 | 1 | 1 | - | 1 |
| 22 | 5 | 8 | 0.8 | 0.125 | 1.200 |
| 23 | 8 | 11 | 0.393 | 0.367 | 1.929 |
| 24 | 3 | 3 | 1 | 0 | 1 |
| 25 | 7 | 9 | 0.429 | 0.444 | 1.762 |
| 26 | 2 | 1 | 1 | - | 1 |
| 27 | 2 | 1 | 1 | - | 1 |
| 28 | 6 | 15 | 1 | 0 | 1 |
| 29 | 2 | 1 | 1 | - | 1 |
| 30 | 2 | 1 | 1 | - | 1 |
| 31 | 2 | 1 | 1 | - | 1 |
| 32 | 2 | 1 | 1 | - | 1 |
| 33 | 2 | 1 | 1 | - | 1 |
| 34 | 2 | 1 | 1 | - | 1 |

Table S4. Summary of the structural properties of the unipartite MAT idiomorph graph considering the whole network. the main subgraph and each individual module. The table includes: a) the number of nodes per graph or subgraph, b) the number of edges, c) the edge density, the ratio of the number of edges and the number of possible edges; d) the average betweenness centrality, which measures the frequency with which the shortest paths between any two nodes passes through each one of them; e) Average path length, which measures the distance between two nodes measured in the number of edges between them.

Table S5. Statistical description of the the unipartite MAT idiomorph graph considering the statistical distribution of descriptors per node for the whole network. the main subgraph and each individual module. The table includes the degree of the network. closeness and betwenness.

| | quantile | degree | closeness | betweenness |
|---|---|---|---|---|
| complete | 2.5% | 1 | 0.196 | 0 |
| | 50% | 15.5 | 0.294 | 0 |
| | 97.5% | 74 | 1 | 0.025 |
| main | 2.5% | 2 | 0.195 | 0 |
| | 50% | 18 | 0.282 | 0 |
| | 97.5% | 74 | 0.379 | 0.033 |
| 1 | 2.5% | 2.9 | 0.417 | 0 |
| | 50% | 42 | 0.678 | 0 |
| | 97.5% | 51 | 0.736 | 0.076 |
| 2 | 2.5% | 1 | 0.346 | 0 |
| | 50% | 10 | 0.643 | 0 |
| | 97.5% | 13 | 0.754 | 0.284 |
| 4 | 2.5% | 1.95 | 0.368 | 0 |
| | 50% | 19 | 0.585 | 0 |
| | 97.5% | 23 | 0.691 | 0.084 |
| 5 | 2.5% | 1.925 | 0.397 | 0 |
| | 50% | 17 | 0.617 | 0 |
| | 97.5% | 28 | 0.804 | 0.108 |
| 6 | 2.5% | 2.15 | 0.373 | 0 |
| | 50% | 39 | 0.700 | 0 |
| | 97.5% | 50 | 0.829 | 0.032 |
| 7 | 2.5% | 3 | 0.235 | 0 |
| | 50% | 6 | 0.299 | 0 |
| | 97.5% | 15 | 0.400 | 0.438 |
| 9 | 2.5% | 3.2 | 0.422 | 0 |
| | 50% | 13 | 0.629 | 0 |
| | 97.5% | 17 | 0.815 | 0.147 |
| 11 | 2.5% | 2 | 0.266 | 0 |
| | 50% | 8 | 0.396 | 0 |
| | 97.5% | 16 | 0.531 | 0.154 |
| 12 | 2.5% | 12 | 0.519 | 0 |
| | 50% | 43 | 0.831 | 0.003 |
| | 97.5% | 44.3 | 0.848 | 0.103 |
| 14 | 2.5% | 4 | 0.269 | 0 |
| | 50% | 9 | 0.446 | 0 |
| | 97.5% | 25.75 | 0.611 | 0.300 |
| 15 | 2.5% | 2.65 | 0.504 | 0 |
| | 50% | 69 | 0.936 | 0 |
| | 97.5% | 70.175 | 0.952 | 0.029 |
| 16 | 2.5% | 5.65 | 0.695 | 0 |
| | 50% | 10 | 0.917 | 0 |
| | 97.5% | 11 | 1 | 0.032 |
| 18 | 2.5% | 2 | 0.256 | 0 |

| | | | | |
|---|---|---|---|---|
| | 50% | 7 | 0.442 | 0 |
| | 97.5% | 22.925 | 0.630 | 0.207 |
| 3 | 2.5% | 1.075 | 0.611 | 0 |
| | 50% | 2 | 0.750 | 0 |
| | 97.5% | 2.925 | 0.981 | 0.616 |
| 8 | 2.5% | 1 | 1 | _ |
| | 50% | 1 | 1 | _ |
| | 97.5% | 1 | 1 | _ |
| 10 | 2.5% | 1 | 1 | _ |
| | 50% | 1 | 1 | _ |
| | 97.5% | 1 | 1 | _ |
| 13 | 2.5% | 3 | 0.615 | 0 |
| | 50% | 5 | 0.727 | 0 |
| | 97.5% | 7.4 | 0.945 | 0.429 |
| 17 | 2.5% | 3 | 1 | 0 |
| | 50% | 3 | 1 | 0 |
| | 97.5% | 3 | 1 | 0 |
| 19 | 2.5% | 4 | 1 | 0 |
| | 50% | 4 | 1 | 0 |
| | 97.5% | 4 | 1 | 0 |
| 20 | 2.5% | 1 | 1 | _ |
| | 50% | 1 | 1 | _ |
| | 97.5% | 1 | 1 | _ |
| 21 | 2.5% | 1 | 1 | _ |
| | 50% | 1 | 1 | _ |
| | 97.5% | 1 | 1 | _ |
| 22 | 2.5% | 2.1 | 0.680 | 0 |
| | 50% | 3 | 0.800 | 0 |
| | 97.5% | 4 | 1 | 0.167 |
| 23 | 2.5% | 1.175 | 0.370 | 0 |
| | 50% | 2.5 | 0.542 | 0 |
| | 97.5% | 4.825 | 0.700 | 0.476 |
| 24 | 2.5% | 2 | 1 | 0 |
| | 50% | 2 | 1 | 0 |
| | 97.5% | 2 | 1 | 0 |
| 25 | 2.5% | 2 | 0.500 | 0 |
| | 50% | 2 | 0.500 | 0 |
| | 97.5% | 4 | 0.750 | 0.533 |
| 26 | 2.5% | 1 | 1 | _ |
| | 50% | 1 | 1 | _ |
| | 97.5% | 1 | 1 | _ |
| 27 | 2.5% | 1 | 1 | _ |
| | 50% | 1 | 1 | _ |
| | 97.5% | 1 | 1 | _ |
| 28 | 2.5% | 5 | 1 | 0 |
| | 50% | 5 | 1 | 0 |
| | 97.5% | 5 | 1 | 0 |
| 29 | 2.5% | 1 | 1 | _ |
| | 50% | 1 | 1 | _ |
| | 97.5% | 1 | 1 | _ |
| 30 | 2.5% | 1 | 1 | _ |
| | 50% | 1 | 1 | _ |
| | 97.5% | 1 | 1 | _ |
| 31 | 2.5% | 1 | 1 | _ |

| | | | | |
|---|---|---|---|---|
| | 50% | 1 | 1 | _ |
| | 97.5% | 1 | 1 | _ |
| 32 | 2.5% | 1 | 1 | _ |
| | 50% | 1 | 1 | _ |
| | 97.5% | 1 | 1 | _ |
| 33 | 2.5% | 1 | 1 | _ |
| | 50% | 1 | 1 | _ |
| | 97.5% | 1 | 1 | _ |
| 34 | 2.5% | 1 | 1 | _ |
| | 50% | 1 | 1 | _ |
| | 97.5% | 1 | 1 | _ |
